# Disease diagnostics using machine learning of immune receptors

**DOI:** 10.1101/2022.04.26.489314

**Authors:** Maxim E. Zaslavsky, Erin Craig, Jackson K. Michuda, Nidhi Sehgal, Nikhil Ram-Mohan, Ji-Yeun Lee, Khoa D. Nguyen, Ramona A. Hoh, Tho D. Pham, Katharina Röltgen, Brandon Lam, Ella S. Parsons, Susan R. Macwana, Wade DeJager, Elizabeth M. Drapeau, Krishna M. Roskin, Charlotte Cunningham-Rundles, M. Anthony Moody, Barton F. Haynes, Jason D. Goldman, James R. Heath, Kari C. Nadeau, Benjamin A. Pinsky, Catherine A. Blish, Scott E. Hensley, Kent Jensen, Everett Meyer, Imelda Balboni, Paul J Utz, Joan T. Merrill, Joel M. Guthridge, Judith A. James, Samuel Yang, Robert Tibshirani, Anshul Kundaje, Scott D. Boyd

## Abstract

Clinical diagnosis typically incorporates physical examination, patient history, and various laboratory tests and imaging studies, but makes limited use of the human system’s own record of antigen exposures encoded by receptors on B cells and T cells. We analyzed immune receptor datasets from 593 individuals to develop *MAchine Learning for Immunological Diagnosis (Mal-ID)*, an interpretive framework to screen for multiple illnesses simultaneously or precisely test for one condition. This approach detects specific infections, autoimmune disorders, vaccine responses, and disease severity differences. Human-interpretable features of the model recapitulate known immune responses to SARS-CoV-2, Influenza, and HIV, highlight antigen-specific receptors, and reveal distinct characteristics of Systemic Lupus Erythematosus and Type-1 Diabetes autoreactivity. This analysis framework has broad potential for scientific and clinical interpretation of human immune responses.

## Main Text

Modern medical diagnosis relies heavily on laboratory testing for cellular or molecular abnormalities, such as the presence of pathogenic microorganisms (*1*, *2*) in the context of clinical history and physical exam findings of infectious disease. For autoimmune diseases such as systemic lupus erythematosus, multiple sclerosis, or type-1 diabetes, there is no single pathogenic agent to detect, and therefore a combination of diagnostic approaches is used to integrate patient history, physical exam findings, results of imaging studies, detection of autoantibodies and other laboratory abnormalities, and exclusion of other conditions. This process can be lengthy and complicated by initial misdiagnoses and ambiguous symptoms (*3–7*). Diagnostic medicine currently makes minimal use of data from the adaptive immune system’s diverse B cell receptors (BCR) and T cell receptors (TCR) that provide antigen specificity to immune responses. In response to pathogens, vaccines, and other stimuli, the repertoires of BCRs and TCRs change in composition by clonal expansion of antigen-specific cells, introduction of additional somatic mutations into BCR genes, and selection processes that further reshape lymphocyte populations. Self-reactive lymphocytes can also clonally proliferate and cause autoimmune diseases or other immunological pathologies. Sequencing of BCRs and TCRs from an individual has the potential to provide a single diagnostic test allowing simultaneous assessment for many infectious, autoimmune, and other immune-mediated diseases (*8–11*).

Receptor repertoire sequencing already contributes to diagnosis and treatment response monitoring in the specialized case of lymphocyte malignancies where the BCR or TCR is a marker of the cancer cells (*12*, *13*). Prior research suggests that BCR sequencing can distinguish between some antibody-mediated pathologies (*14*). Challenges to broader application of these methods include low frequencies of antigen-specific B cells and T cells in many patients, the high diversity of immune receptor genes produced by gene rearrangement during lymphocyte development, and somatic hypermutations that accumulate in BCRs following B cell stimulation (*9*, *15*), as well as technical factors including experimental protocols for sequence library preparation, and differences in patient demographics or past exposures that may influence the responses to a given antigen (*16*). Previous investigations of disease or vaccination-related immune repertoires have identified with varying degrees of success similar receptor amino acid sequences, subsequences, or somatic mutation patterns in people with the same exposures (*17–28*). Others have represented receptor sequences with alternative encodings of amino acid biochemical properties such as charge and polarity to improve detection of receptor groups of similar antigen specificity (*29–32*). Learned representations of BCRs or TCRs from language models and variational autoencoders are candidates for immune state classification and for predictive applications such as therapeutic antibody optimization (*33–42*). Probabilistic models of V(D)J recombination and selection processes have also been applied to better understand immune receptor generation and expansion in response to antigenic stimuli (*43*, *44*). Very few studies have attempted to integrate BCR and TCR data for diagnostic purposes, however, and it remains unclear to what extent immune repertoire sequence data are sufficient for generalized and accurate infectious or immunological disease classification.

To address these challenges, we have developed *MAchine Learning for Immunological Diagnosis (Mal-ID),* which combines three machine learning representations for both BCR and TCR repertoires to detect infectious or immunological diseases in patients (**Fig. 1**). *Mal-ID* uses overview summary metrics of receptor populations and focused analysis of the key CDR3 (complementarity region 3) antigen-binding loop with sequence distance measures and protein language modeling. We applied *Mal-ID* to 16.2 million BCR heavy chain clones and 23.5 million TCR beta chain clones systematically collected from peripheral blood samples of 593 individuals with different immunological statuses, as well as external datasets collected with different experimental protocols. Without prior knowledge of pathogenesis or of which sequences are antigen specific, *Mal-ID* distinguishes healthy from sick individuals, viral infections from autoimmune conditions, different infections from each other, and different autoimmune diseases from each other. The results support the robustness of immune receptor-based diagnosis for identifying diverse immune states and show correlation with autoimmune disease severity. Importantly, the model learns to prioritize antigen specific sequences for patient diagnosis.

**Fig. 1.**
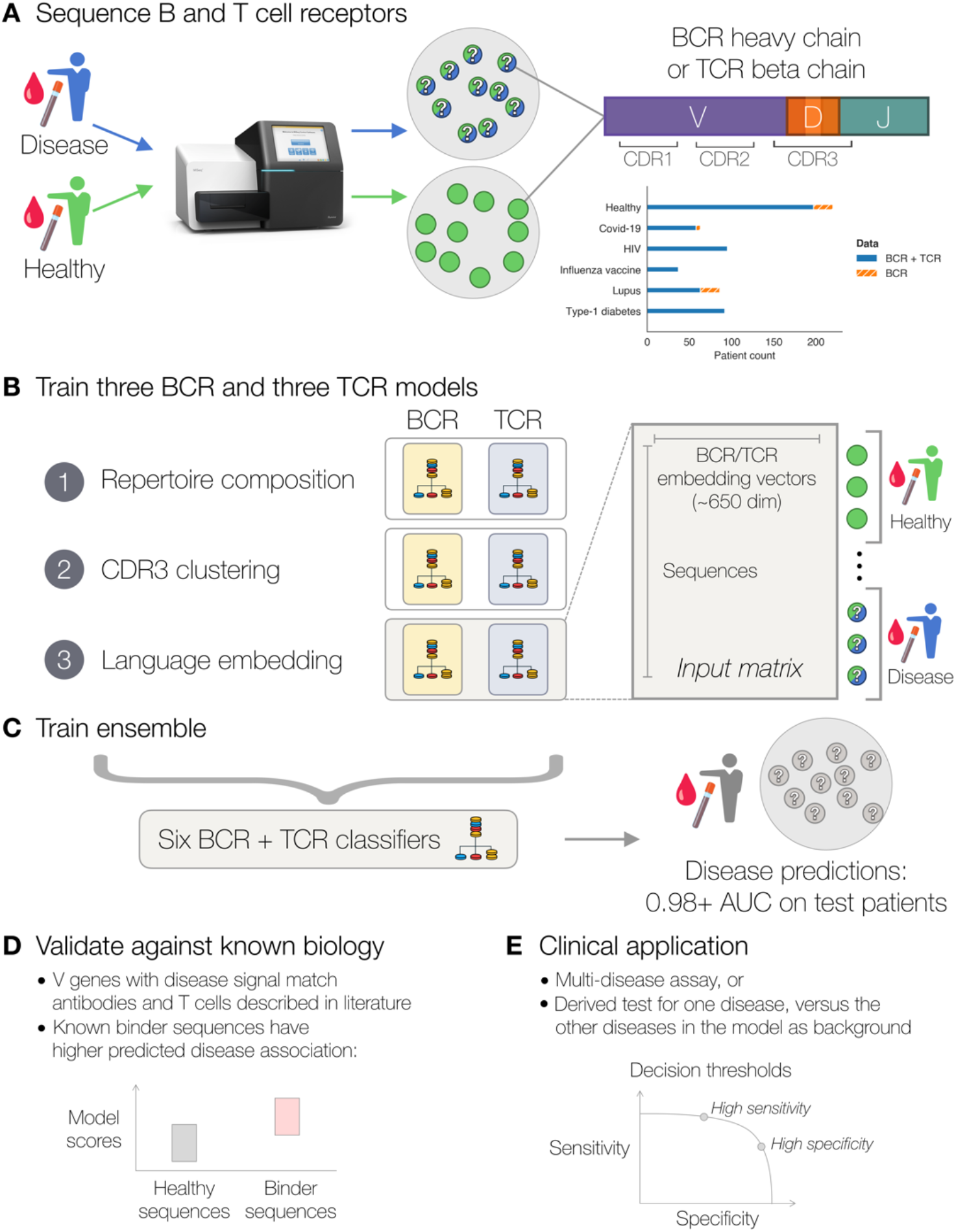
MAchine Learning for Immunological Diagnosis (*Mal-ID*) framework. **(A)** BCR heavy chain and TCR beta chain gene repertoires are amplified and sequenced from blood samples of individuals with different disease states. Question marks indicate that most sequences from patients are not disease specific. **(B)** Machine learning models are trained to predict disease using several immune repertoire feature representations. These include protein language models, which convert each amino acid sequence into a numerical vector. **(C)** An ensemble disease predictor is trained using the three BCR and three TCR base models. The combined model predicts disease status of held-out test individuals. **(D)** For validation, the disease prediction model allows introspection of which V genes carry disease-specific signal, which can be validated against prior literature. Within each V gene, previously published BCR and TCR sequences known to be disease associated can be tested for whether they have higher disease association. **(E)** The final trained model can be applied as a multi-disease assay, or as a diagnostic test for one disease. The same model will achieve a range of sensitivities and specificities depending on the chosen decision threshold.

### Integrated repertoire models of immune states

*Mal-ID* uses three models per gene locus (BCR heavy chain, IgH; and TCR beta chain, TRB) to recognize immune states (**Fig. 1B, fig. S1**). Each model focuses on different aspects of immune repertoires shared between individuals with the same immune state or diagnosis: gene segment frequencies and IgH mutation rates in each isotype (Model 1), highly similar CDR3 sequence clusters (Model 2), and inferred potential structural or binding similarity based on ESM-2 protein language model embeddings of CDR3 sequences (*45*) (Model 3). Outputs from the three BCR and three TCR models are combined into a final prediction of immune status with a logistic regression ensemble model that can resolve potential errors of individual predictors (*46*). The trained program takes an individual’s peripheral blood BCRs and TCRs as input and predicts the probability of each disease on record (**Fig. 1C**). Full details of the modeling approach are provided in **Materials and Methods**.

We applied this approach to patients diagnosed with Covid-19 (n=63), HIV infection (n=95) (*18*), Systemic Lupus Erythematosus (SLE, n=86), and Type-1 Diabetes (T1D, n=92), as well as influenza vaccination recipients (n=37) and healthy controls (n=220) (**table S1**). All datasets used a standardized sequencing protocol to minimize batch effects (**Materials and Methods**). To evaluate generalizability, patients were strictly separated into training, validation, and testing sets (**fig. S2**). Any repeated samples from the same individual were kept grouped together during this division process, to ensure data from the same individual did not leak between training and testing steps. We trained separate models per cross-validation fold and report averaged classification performance. As described below, we further tested for the potential contribution of batch effects and demographic differences to diagnostic accuracy.

The ensemble approach distinguished six specific disease states in 550 paired BCR and TCR samples from 542 individuals with a multi-class area under the Receiver Operating Characteristic curve (AUROC) score of 0.986 (**Fig. 2A**). AUROC is the likelihood of correctly ranking positive examples higher than negative examples (*47*), averaged across all disease label pairs weighted by frequency. Other performance metrics are provided in **table S2**.

**Fig. 2.**
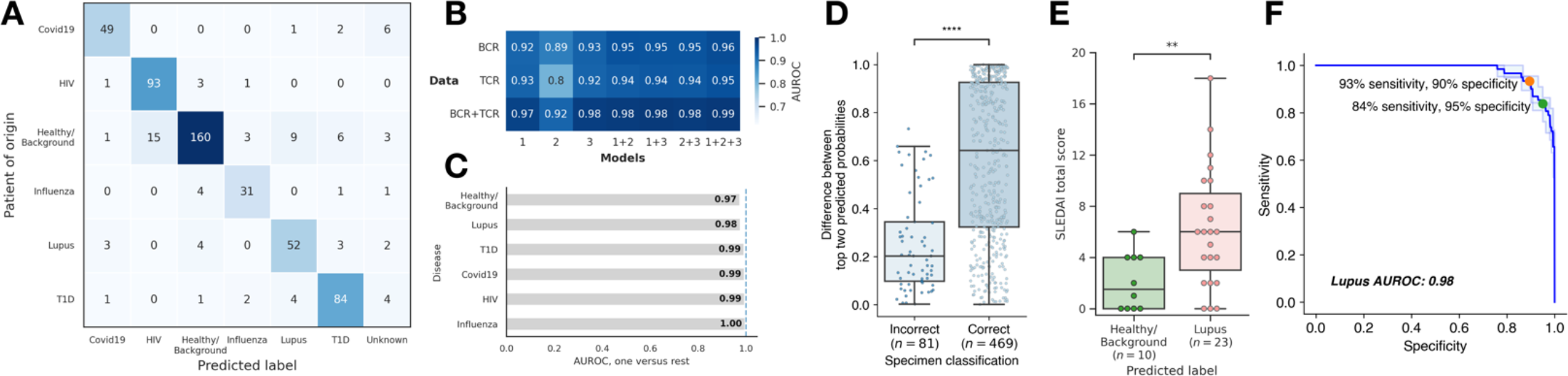
*Mal-ID* classifies disease using IgH and TRB sequences. (**A**) Disease classification performance on held-out test data by the ensemble of three B cell repertoire and three T cell repertoire machine learning models, combined over all cross-validation folds. (**B**) Disease classification performance (multi-class one-vs-one AUROC scores) divided column-wise by model architecture (individual base models or ensembles of base models) and row-wise by whether BCR data, TCR data, or both were incorporated. Model 1 refers to the repertoire composition classifier, model 2 refers to the CDR3 clustering classifier, and model 3 refers to the protein language model classifier. The CDR3 clustering models abstain from prediction on some samples, while the other models do not abstain; to make the scores comparable, abstentions were forcibly applied to the other models. The BCR-only results also include BCR-only patient cohorts (n=66 samples) not present in TCR-only or BCR+TCR evaluation. (**C**) AUROC scores for each class versus the rest from the full ensemble architecture including models 1, 2, and 3 with both BCR and TCR data. (**D**) Difference of probabilities of the top two predicted classes for correct versus incorrect ensemble model predictions. A higher difference implies that the model is more certain in its decision to predict the winning disease label, whereas a low difference suggests that the top two possible predictions were a toss-up. Results were combined across all cross-validation folds. One-sided Wilcoxon rank-sum test: p value 1.599 × 10^−15^, U-statistic 6052. (**E**) SLEDAI clinical disease activity scores for adult lupus patients who were either classified correctly or misclassified as healthy by the BCR-only ensemble model, used here because the adult lupus data was primarily BCR-only. SLEDAI scores were only available for some patients. One-sided Wilcoxon rank-sum test: p value 4.242 × 10^−3^, U-statistic 48. (**F**) Sensitivity versus specificity, averaged over three cross-validation folds, for a lupus diagnostic classifier derived from the pan-disease classifier. Two possible decision thresholds are highlighted.

*Mal-ID* outperforms previously reported classification approaches on our evaluation dataset. The CDR3 clustering model, similar to convergent or public sequence discovery approaches in the literature, achieves only 0.89 AUROC for BCR and 0.80 AUROC for TCR (**Fig. 2B**). Another approach based on exact sequence matches, originally reported for TCR sequences (*17*), achieves 41% accuracy for BCR data and finds no hits in 40% of samples (**fig. S3**). Identical sequences across individuals are expected to be rare for IgH because of somatic hypermutation, but the HIV class was an exception. For TCR data, the exact matches technique almost always finds hits, but achieves only 42% accuracy and 0.75 AUROC and predicts that almost all samples belong to either the Covid-19 class or the healthy class. *Mal-ID*’s AUROC of over 0.98 represents a major increase in diagnostic accuracy.

The three-model approach discriminates between autoimmune diseases, viral infections, and influenza vaccine recipient samples collected at day seven after vaccination, when B cells responding to the vaccine are usually at peak frequencies (*48*, *49*). The different BCR and TCR components of the ensemble model contributed to varying degrees for classification of each immunological condition (**Fig. 2B, fig. S4**). TCR sequencing provided more relevant information for lupus and type-1 diabetes, while Covid-19, HIV, and influenza had clearer BCR signatures. Combined BCR and TCR data performed best (**table S2**). Alone, the repertoire composition Model 1 and protein language embedding Model 3 classifiers performed better on average than the CDR3 clustering Model 2; of these, the TCR CDR3 clustering model was the weakest, potentially because the model did not account for patient HLA genotypes. The combination of Models 1, 2, and 3 generally had best performance, but pairing Models 1 and 3 performed as well for many classes (**fig. S4**), suggesting CDR3 clustering may not be required for classification or is encompassed by the protein language model results.

In practice, decision thresholds to categorize patient samples into disease categories can be chosen depending on the consequences of different types of errors, the performance metrics to be optimized, and the priority given to different diseases. Here, we illustrate how the estimated AUROCs translate to explicit misclassification rates for a few different case studies. Using a naïve assignment scheme, where we assign each patient to the class with the highest predicted probability, *Mal-ID* achieves 85.3% accuracy (**Fig. 2A**). Among misclassified repertoires, 2.9% lacked sequences belonging to Model 2 CDR3 clusters, making the CDR3 clustering component abstain from prediction. The remaining 11.8% had inconclusive predictions (**Fig. 2D**). Many misclassifications involved healthy donors predicted as having an illness, indicating that the model selecting classification labels based on the highest prediction probabilities resulted in more false positive than false negative results. Some of these errors may also be caused by healthy control individuals not being screened for definitive absence of all the diseases in our panel. However, 92.9% of sick patients and vaccine recipients were identified as not being in a healthy/baseline immune state, and 87.5% had their particular immune state properly classified. Adult lupus patients were the most challenging disease category to classify (**Fig. 2C**). Unlike the pediatric lupus cohort, the adults were on therapy, which can influence immune repertoires (*14*). Most adult lupus patients had BCR data only. Based on this more limited data, a subset of patients was predicted as healthy (**fig. S5**). However, misclassified patients had significantly lower Systemic Lupus Erythematosus Disease Activity Index (SLEDAI) scores (*50*, *51*) (**Fig. 2E**), indicating better-controlled or quiescent disease in response to treatment, which likely influenced the model’s tendency to classify them as immunologically healthy. Compared to the 85.3% overall accuracy achieved by the model using BCR and TCR data together, the BCR-only and TCR-only versions of *Mal-ID* had 74.0% and 75.1% accuracy (**table S2**), respectively, further highlighting the benefit of analyzing BCR and TCR data jointly when it is available.

Disease-specific classifiers can also be trained or derived from the pan-disease model. For example, by labeling lupus predictions as positives and others as negatives, we extracted a lupus diagnosis model, which is clinically relevant due to the lack of a sensitive and specific lupus test (*52*, *53*). Adjusting the decision threshold for high lupus sensitivity, our model achieves 97% sensitivity and 86% specificity, or 84% sensitivity and 95% specificity when optimized for specificity (**Fig. 2F**). Balanced performance of 93% sensitivity and 90% specificity is also possible. Therefore, *Mal-ID* can serve as a multi-disease test or be specialized for detecting a particular condition.

### Limited impact of batch effects on classification

To assess *Mal-ID*’s generalizability, we trained a model on all the data (**fig. S2**), then tested on Covid-19 patient and healthy donor repertoires from other BCR or TCR studies with similar complementary DNA (cDNA) sequencing protocols. *Mal-ID* predicted disease in two BCR external cohorts (*54*, *55*) with perfect 1.0 AUROC: all seven Covid-19 patients received higher Covid-19 predicted probabilities than did the six healthy donors. However, accuracy was 69% by naïve assignment: one Covid-19 patient was misclassified as type-1 diabetes, and three healthy donors were misclassified as lupus or type-1 diabetes (**fig. S6A**, **table S3**). As the base rates of disease have changed in this evaluation dataset containing only Covid-19 patients and healthy donors, the decision thresholds should be tuned; the adjustment can be learned automatically by holding out a small portion of the external cohorts. After this tuning, the adjusted BCR model reached 100% accuracy in the remaining evaluation data (**fig. S6B**). Similar tuning could be performed for clinical contexts with varying disease prevalence.

In TCR external cohorts of 17 Covid-19 patients and 39 healthy donors (*56–58*), *Mal-ID* achieved 0.99 AUROC and 68% accuracy based on naïve assignment, which rose to 90% accuracy after threshold tuning (**fig. S6, C** and **D, table S3**). Almost all Covid-19 patients and healthy donors evaluated (i.e. not held out for tuning) were correctly identified, except 3 of 28 healthy donors were misclassified as Covid-19, and 1 of 12 Covid-19 patients was misclassified as healthy. Low accuracy prior to tuning was caused by misclassifications of Covid-19 patients as lupus due to Model 2, which also performed poorly on our primary TCR data as noted above. Disabling Model 2 led to no Covid-19 patients misclassified as lupus and 89% accuracy without tuning (along with 0.97 AUROC). High performance on new datasets suggests that *Mal-ID* learns generalizable disease-related signals and is robust even with only BCR or only TCR data. This adaptability extends to different sequencing modalities, including TCR genomic DNA sequencing from Adaptive Biotechnologies. Observing gene segment usage distinct from cDNA data as previously reported (*16*) (**fig. S7A**), we retrained *Mal-ID* to successfully separate six immune states in 1365 samples: common variable immunodeficiency (CVID), Covid-19, HIV, rheumatoid arthritis (RA), T1D, and healthy. These studies were conducted by different labs, introducing the possibility of batch effects, and are restricted to only TCR data (**table S4**). *Mal-ID* classifies these disease classes with 0.97 AUROC and 88% accuracy (**fig. S7B**), indicating *Mal-ID* can learn disease signals across sequencing modalities and scales to over 150 million sequences. As in the primary *Mal-ID* dataset, misclassifications often involved healthy individuals being predicted as sick, but 96% of sick patients were correctly identified as having an illness. The Covid-19 and healthy data came from studies that were divided into multiple cohorts; for example, the Emerson *et al.*, 2017 study of healthy individuals included an original cohort and an independent validation cohort (*17*). Therefore, we also trained *Mal-ID* with these cohort divisions preserved. Holding out entire Covid-19 and healthy cohorts from the training process, we saw that *Mal-ID* accurately classified the independent cohorts with 1.0 AUROC and 98% accuracy (**fig. S7C**).

To test for batch effects in our primary data, we retrained *Mal-ID* holding out an entire Covid-19 cohort of 10 patients, as well as 13 healthy samples that were resequenced in a new replicate batch. All held-out Covid-19 samples and healthy samples (pooling the original and replicate data) were correctly classified. When we split each healthy donor’s replicates, both replicates were correctly classified for 9 of 13 healthy donors with 97% or higher correlation between predicted class probabilities, while two individuals had replicates with abstention from classification, and two had divergent classification for each replicate (**fig. S8**). Classification abstentions resulted from two replicates matching no class-associated CDR3 clusters, which was likely caused by these replicates having fewer IgH clones than the rest due to limited sequencing depth. The accurate classification of a new Covid-19 cohort and consistent scoring of healthy replicate samples increases the likelihood that *Mal-ID* learns true biological signal rather than batch effects.

### Limited impact of age, sex, and race on classification

Patient demographics also influence the immune repertoire (*57–60*). To evaluate how extraneous covariates may affect classification, we attempted to predict age, sex, or ancestry from the immune repertoires of healthy individuals. While sex could not be accurately determined, sequences carried relatively weak ancestry signals (0.78 AUROC, **table S5**). Ancestry separation is visible in gene segment usage (**fig. S9A**), potentially from germline IgH and TRB locus differences, shaping of TCR repertoires by HLA alleles that differ between ancestry groups, and different environmental exposures in the African ancestry individuals living in Africa in the data (*61*, *62*). Consistent with potential influences of HLA genotype, *Mal-ID*’s TCR components had less accuracy in distinguishing HIV patients and healthy controls from the African cohort. The corresponding IgH repertoires were more distinct (**fig. S10**), highlighting the advantage of combining BCR and TCR data.

Previous studies noted age-related changes in gene expression, cytokine levels, and immune cell frequencies (*63*, *64*). We observed a modest age signal in healthy IgH and TRB sequences, achieving 0.75 AUROC for distinguishing age 50 and up, excluding 19% of samples that matched no CDR3 age clusters (54% accuracy including abstentions; **table S5**). Age signatures may correspond to imprinting effects from childhood exposure to viruses such as influenza (*65*) or to autoreactivity increasing with age (*66*). Pediatric samples had especially distinct TRBV gene usage (**fig. S9B**), and *Mal-ID* identified them with perfect 1.0 AUROC when it made predictions (**table S5**), though accuracy was 55% due to 45% abstention. Despite substantial differences in the remaining samples, age effects did not interfere with disease classification: *Mal-ID* accurately distinguished pediatric patients and controls (**fig. S5C**). The high Model 2 abstention rates indicate relatively few age-associated CDR3 sequence clusters, and show that unsupervised clustering will not necessarily choose clusters that correspond to age or other desired axis of variation. Also, we restricted *Mal-ID*’s scope to B cell populations shaped by antigenic stimulation: somatically hypermutated IgD/IgM and class switched IgG/IgA isotypes. Studying naive B cells may reveal additional age, sex, or ancestry effects.

To assess whether demographic differences between disease cohorts drove our classification results, we attempted to predict disease state from age, sex, and ancestry alone, ignoring sequence data. Ages by cohort were: T1D median 14.5 years (range 2-74); SLE median 18 years (range 7-71); influenza vaccine recipient median 26 years (range 21-74); HIV median 31 years (range 19-64); healthy control median 34.5 years (range 8-81); Covid-19 median 48 years (range 21-88) (**table S1**). The percentage of females in each cohort was 50% (healthy controls), 52% (Covid-19), 57% (influenza vaccine recipient), 64% (HIV), and 85% (SLE), consistent with high representation of females in SLE (*67*). The ancestries and geographical locations of participants also differed between cohorts. Notably, 89% of individuals in the HIV cohort lived in Africa (*18*). Using only age, sex, or ancestry, disease AUROCs were 0.68, 0.59, and 0.79, respectively. A classifier with all three features achieved 0.85 AUROC, substantially lower than the 0.98 AUROC from *Mal-ID* retrained with demographics alongside sequence features (**table S6**, **fig. S11, A** and **B**). To underscore the significant disease signal in BCRs and TCRs, we also evaluated the demographics-only classifier on the external cDNA datasets. For TCR, it achieved 0.48 AUROC and 50% accuracy (**fig. S6F**), compared to *Mal-ID*’s 0.99 AUROC and 68% accuracy before threshold tuning (**table S3**). For BCR, the demographics-only classifier achieved 1.0 AUROC identical to the standard *Mal-ID* model because the external Covid-19 patients were all Asian, while the healthy controls were Caucasian or African American. Nevertheless, accuracy was 58% with demographic features (**fig. S6E**), compared to 69% with *Mal-ID* before tuning (**table S3**). Demographic covariates therefore do not explain model performance on external validation data. As an additional test to confirm that predictions were not driven by demographics, we retrained with age, sex, and ancestry effects regressed out from the ensemble model’s feature matrix. Classification performance for individuals with known demographics dropped slightly from 0.98 AUROC to 0.96 AUROC after decorrelating sequence features from demographic covariates (**table S6**, **fig. S11C**), suggesting age, sex, and ancestry have modest impacts on disease classification.

### Language model recapitulates immunological knowledge

To better understand the factors contributing to the high accuracy of *Mal-ID* classification, we asked which biological patterns identified each disease. Model 3 reveals which receptor sequences contribute most to disease predictions because BCRs or TCRs are scored individually, then aggregated into patient predictions. Separate models generate sequence predictions specialized for each IGHV gene and isotype combination in the BCR case, or for each TRBV gene in TCR data (**Materials and Methods**). We calculated Shapley importance (SHAP) values (*68*, *69*) for the disease probabilities derived from each sequence category, which serve as features for making Model 3’s patient predictions. V genes and isotypes are given priority in the aggregation model based on their prevalence in patients and on containing sequences distinct from other immune states by CDR3 features. According to V gene category contributions to disease predictions, our model’s classifications align with established immunological knowledge from data such as antigen-specific B cell and T cell isolation and receptor sequencing (**Supplementary Text**). For example, IGHV1-24 and IGHV2-70 are prioritized for Covid-19 prediction, IGHV4-34 and IGHV4-59 have greater weight for lupus, IGHV1-2 and IGHV4-34 for HIV, and IGHV3-23 for influenza (**Fig. 3**). We also decomposed the lupus and T1D SHAP values into TRBV gene prioritization clusters corresponding to patient age (**figs. S12-13**). In our lupus cohort, age is associated with treatment status, as the adults were on treatment while the pediatric cohort was treatment naive, indicating that differences in gene usage may also depend on treatment.

**Fig. 3:**
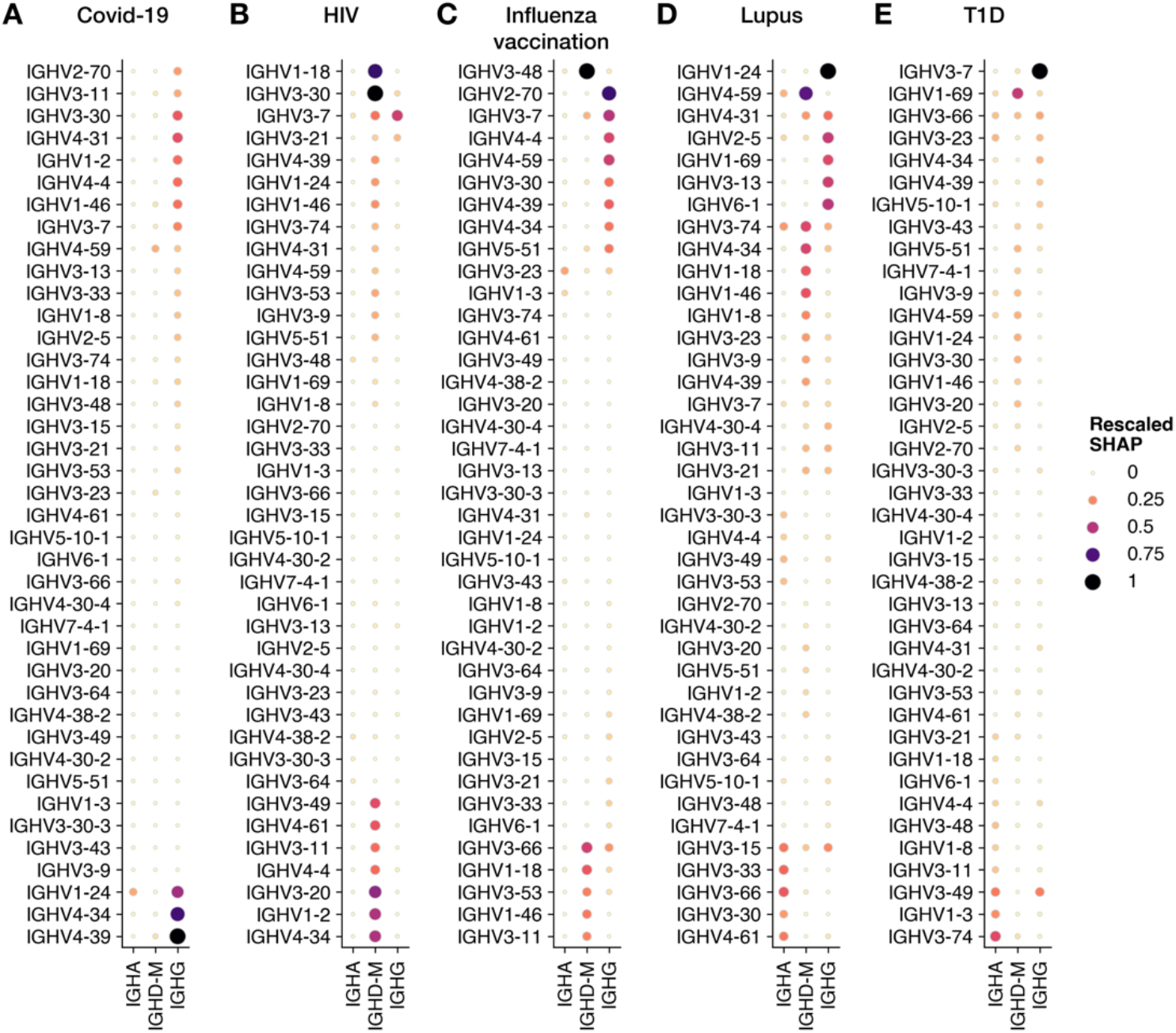
Disease-associated IGHV genes and isotypes prioritized by Model 3 using protein language embeddings. Shapley importance (SHAP) values quantifying the contribution of average sequence predictions from each IGHV gene and isotype category to Model 3’s prediction of a sample’s disease state are plotted for (**A**) Covid-19, (**B**) HIV, (**C**) influenza vaccination, (**D**) lupus, and (**E**) type-1 diabetes.

Different diseases showed varied association of IGHV gene usage in the context of particular BCR heavy chain isotypes. Covid-19 prediction prioritized IgG (**Fig. 3A**), as expected from prominent IgG expression by SARS-CoV-2-specific B cells (*70–73*). While IgA contributions were minimal for Covid-19, HIV, and influenza predictions, IgA was informative for lupus, consistent with disease-associated IgA autoantibodies described in the literature (*74*), as well as for T1D, along with other isotypes (**Fig. 3, D** and **E**). The HIV model favored mutated IgM/D (**Fig. 3B**). Influenza predictions were driven by IgG and mutated IgM/D signal primarily (**Fig. 3C**). B cell isotype usage varied by person and across disease cohorts (**fig. S14**), but the model also considers distinct disease signal enrichment within each isotype to determine its priority. Other *Mal-ID* components were not influenced by isotype sampling variation: Model 1 quantifies each isotype group separately, and Model 2 is blind to isotype information. To be sure that differences in isotype proportions between patient cohorts were insufficient to predict disease, we attempted to predict disease from a sample’s isotype proportions without any sequence information, achieving only 0.68 AUROC compared to *Mal-ID*’s AUROC of over 0.98.

Having validated that V gene segments and isotypes prioritizations for disease identification match the literature, we assessed whether the multi-disease *Mal-ID* model could distinguish reported SARS-CoV-2 binding BCRs (*75*) from healthy donor sequences, despite having been trained for patient rather than sequence classification (**Supplementary Text**). Model 3 assigned higher Covid-19 probabilities to reported binders compared to healthy sequences for IGHV1-24, IGHV2-70, and other key V genes, with AUROC ranging up to 0.78 across IGHV genes and area under the precision-recall curve (AUPRC) up to 6.9-fold over baseline (**Fig. 4, E** to **G**). Model 2 Covid-19 associated clusters identified some known binders, with up to 100% precision in IGHV1-24 and IGHV3-53 among others, but low recall (**Fig. 4, A** to **D**). The higher ranking of experimentally validated, disease-specific sequences from separate cohorts suggests that the models learn antigen-specific sequence patterns within important IGHV genes that recapitulate biological knowledge gained during the extraordinary international research effort in response to the Covid-19 pandemic, despite the enormous diversity of immune receptor sequences, and despite being trained without knowledge of which Covid-19 patient BCRs were specific for SARS-CoV-2 antigens. Only a fraction of peripheral blood B and T cell receptor sequences from Covid-19 patients are thought to be directly related to the SARS-CoV-2 viral antigen-specific immune response (*76*, *77*). However, cDNA sequencing may emphasize plasmablasts with high RNA copy counts, and excluding naive B cells may highlight antigen-experienced B cells during training.

**Fig. 4.**
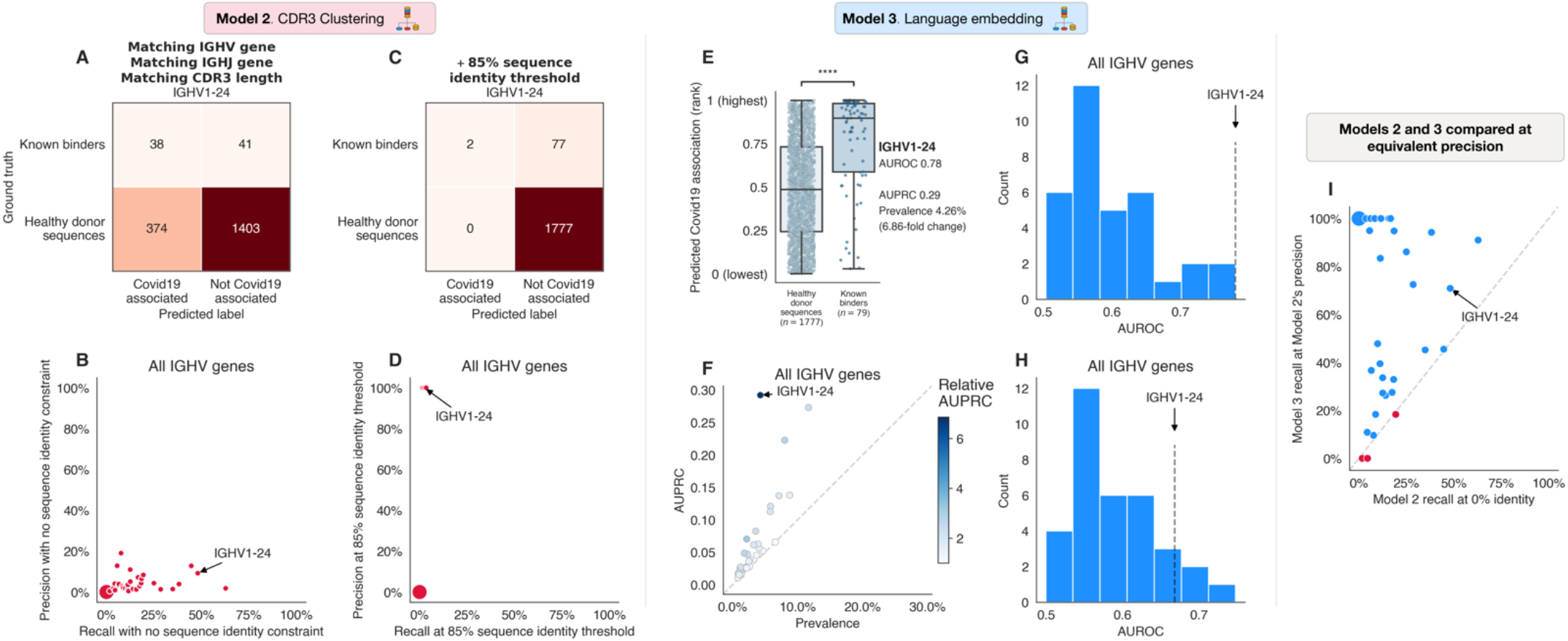
Models 2 and 3 learn SARS-CoV-2 antigen-specific sequence patterns from Covid-19 patient data and can distinguish between known SARS-CoV-2-specific antibody sequences and healthy donor sequences. For this comparison, validated SARS-CoV-2-binding sequences from the CoV-AbDab database (*75*) and a subset of healthy donor sequences were held out from training. Known binder detection using Model 2 or Model 3 predictions of sequence association to disease was evaluated separately for each IGHV gene; performance is shown for IGHV1-24 and compared across IGHV genes. (**A** to **D**) Model 2 identifies a conservative set of public clones enriched in Covid-19 patients which match some known binders. Model 2’s precision and recall across IGHV genes is shown, with binding predictions determined: (**A** and **B**) based on shared IGHV gene, IGHJ gene, and CDR3 length with any Covid-19 cluster identified in Model 2’s training procedure; or (**C** and **D**) at an 85% CDR3 sequence identity threshold. (**E** to **H**) Model 3 ranks known binders higher than healthy sequences based on predicted Covid-19 probability (**E**), with relative AUPRC ranging up to 6.9-fold over baseline prevalence (**F**) and AUROC up to 0.78 across IGHV genes (**G**). Permutation test in panel (E) to assess whether IGHV1-24 known binders have higher ranks than healthy donor sequences, with consistent labels maintained during the permutation process across sequences from each healthy donor: p value 0. (**H**) Model 3 maintains reasonable performance (AUROC up to 0.75) for sequences that are not evaluated by Model 2’s clustering (sequences for which Model 2 identified no SARS-CoV-2 clusters with matching IGHV gene, IGHJ gene, and CDR3 length). (**I**) At equivalent precision, Model 3 generally exhibits higher recall than Model 2, identifying more true binders but with increased false positives. IGHV genes where Model 3 has higher recall than Model 2 are shown in blue. For each IGHV gene, recall was calculated for Models 2 and 3 at Model 2’s precision shown in (B). Point size indicates number of identical values plotted at a particular location for panels (B), (D), and (I).

We repeated the test with influenza known binders (*78*), finding that both models again prioritized binding sequences in key IGHV genes (**Supplementary Text**). However, enrichment was more muted, ranging up to 0.65 AUROC and 4.0-fold change over baseline AUPRC for Model 3. The relatively lower scores may be because the reference influenza-specific antibodies were derived from studies using a small sampling of all the influenza antigens that have been reported over past decades, and were not derived from responses to the annual vaccine of the same year as the samples analyzed in our study. Differences in response to flu infection versus vaccination may also contribute to the relatively lower known binder enrichment scores: unlike the Covid-19 case where the models were trained with data from patients, our influenza training data was limited to vaccinated individuals while the known binders studied were derived from both infected and vaccinated individuals.

Finally, evaluating on SARS-CoV-2-specific TCRs (*79*), Model 2 performed poorly, consistent with the relatively low Model 2 TCR patient classification performance described earlier, while Model 3 scores had weak enrichment for known binders, up to 0.56 AUROC and 1.30-fold AUPRC change in any TRBV gene (**Supplementary Text**). Compared to IgH, TRB known binders may have less enrichment for higher Model 3 ranks over healthy sequences because the interactions between TCR and genetically diverse HLA molecules during T cell stimulation introduce additional differences between cohorts and between participants within cohorts, and activation of T cells upon peptide stimulation in culture may result in some non-specific bystander clone activation. Further, unlike the IgH classification, the TCR analyses do not exclude naive T cells that could contain low frequencies of SARS-CoV-2 specific clones in unexposed individuals. This moderate performance for antigen-specific sequence identification nevertheless leads to high patient diagnosis performance via aggregating many weak sequence-level scores, as has been demonstrated in other contexts where weak learners are ensembled into a strong classifier (*80*). Also, to produce patient diagnosis predictions from TCR data, sequence-level predictions were aggregated simply by calculating average predicted probabilities after filtering out a percentage of low information content sequences (**Materials and Methods**). The strength of the patient predictions achieved by averaging many sequences indicates that diseases may alter immune repertoires by affecting a larger proportion of clones than those that explicitly bind antigens from the stimulus. Therefore, another possible explanation for the moderate enrichment in predicted probabilities for SARS-CoV-2 binding TCRs over healthy TCRs is that the classifier may have learned additional patterns other than those of TCRs that directly bind to the virus.

## Discussion

In this study, we asked whether immune receptor sequencing can accurately determine a person’s disease or immune response state, based on pathogenic exposures and autoreactivity shaping the immune system’s collection of antigen-specific adaptive immune receptors. The three-part machine learning analysis framework we applied to well-characterized datasets of six distinct immunological states classified immune responses with performance of 0.986 AUROC, leveraging both B and T cell signals in 542 individuals. We ensured models were never trained on data from a patient and then evaluated on other data from the same person. Faced with highly diverse repertoires containing tens to hundreds of thousands of unique sequences, the *Mal-ID* ensemble of classifiers learned disease-specific patterns and prioritized meaningful sequences for prediction of specific viral infections and autoimmune diseases. These signatures of specific disease types overrode more modest differences detectable between individuals differing by sex, age, or ancestry. *Mal-ID* generalizes to sequencing data from other laboratories and experimental protocols after additional tuning. Our architecture scales to population-level data; in this study, we demonstrated its use for over 1350 samples at a time with external datasets.

Key innovations for *Mal-ID*’s performance are the trio of analysis models to extract signal from B and T cell repertoires, as well as the way they are combined, fusing aggregate repertoire composition properties, detection of important sequence groups, and language model interpretations of individual sequences. The components are additive: integrating these models outperforms them individually and suggests that they capture different patterns. Combining BCR and TCR repertoire data provides more accurate classification than either receptor type alone, potentially reflecting variation in the roles of B cell and T cell responses in different diseases. For example, type-1 diabetes is considered to be predominantly T cell mediated (*81–83*), and our T cell-only model indeed distinguishes T1D from other classes better than our B cell-only model (0.96 AUROC and 93% accuracy, compared to 0.93 AUROC and 86% accuracy). But by combining both signals, T1D detection performance increases to 0.99 AUROC and 96% accuracy (**fig. S4**). Similarly, lupus can be classified with 0.93-0.94 AUROC and 91% accuracy by either B or T cell information alone, which is supported by the prominence of autoantibodies in this condition (*84*) and the known contributions of T cells to the pathology of SLE (*85*), but it is best classified by the combination of B cell and T cell models, which together achieve 0.98 AUROC and 95% accuracy. These results confirm that B and T cell information considered together in immune response analysis provides a more complete description of the immune state.

The CDR3 clustering and language model components of our model assess which receptor sequences have highest predicted disease association. Sequences independently validated to be pathogen associated were distinguished from healthy donor sequences in the Covid-19 and influenza analyses, confirming that *Mal-ID* learns receptor sequence patterns used in the immune response to disease and vaccination. Labels on individual sequences are not required to train these models. Additionally, the model architecture reveals which sequence categories contribute most to predictions of each disease — which V genes and isotypes are important building blocks for the BCRs and TCRs deployed by the immune system. We confirmed that V genes reported in prior literature carry high weight in the *Mal-ID* prediction process, and our analysis highlighted several V genes as characteristic ones in particular disease conditions that are not yet described in the literature, posing hypotheses that can be tested in future research (**Supplementary Text**). Unlike comparisons limited to patients with one disease versus healthy individuals, which may flag generic inflammatory responses, the multi-class modeling approach in this study can pinpoint immune behaviors specific to each disease type.

With appropriate clinical validation, a model trained with the *Mal-ID* framework could be deployed either as an assay to distinguish several infectious and autoimmune diseases simultaneously, or as a diagnostic test for one particular disease. The trained proof of concept *Mal-ID* model achieves high sensitivity and specificity in distinguishing lupus from another autoimmune disease, several infectious diseases, and healthy controls. Furthermore, our model seems to distinguish lupus patients with milder symptoms from those with significant disease activity. The model can also detect immune responses due to medical interventions such as vaccination. Therefore, this repertoire sequencing and analysis technique has potential value for autoimmune diagnostics and immune monitoring in order to optimize treatments.

In this study, we emphasized empirical data from real patients to produce a large cohort with consistently collected IgH and TRB immune receptor sequencing data. Larger datasets could be generated by simulation, but we avoided this path due to uncertainty associated with selecting features to accurately simulate real data when generating artificial repertoires for method testing. Real data come with potential concerns about batch effects that we attempt to address in several ways. First, we used consistent sample and receptor sequence library preparation, sequencing, and bioinformatics protocols for all data in this study. Second, we trained an alternative model based on demographic covariates only, which showed that demographics do not explain the data adequately and that our primary model performs significantly better. We also demonstrated that a joint model with sequence information and demographic covariates does not perform much better than our primary sequence model. Third, we held out an entire Covid-19 patient cohort, confirming they are properly classified in a validation step. We also performed replicate sequencing for a set of healthy control individuals; holding out these healthy controls from training, we confirmed that most replicates have consistent predictions. Finally, we evaluated performance on completely independent cohorts from other centers; a model that latches onto batch effects would not work in that setting, but *Mal-ID* does. These results indicate that the *Mal-ID* model generalizes to new data and does not fit to latent, unknown hidden variables.

The *Mal-ID* framework appears to capture fundamental principles of immune responses, and it appears to generalize to separate clinical cohorts. The task of differentiating Covid-19, HIV infection, lupus, type-1 diabetes, and healthy was employed as a demonstration of the methodology’s potential. Additional testing will be needed to extend the framework to particular clinical scenarios, and could involve calibrating classification probabilities to generate disease scores with known diagnostic uncertainty, addressing multiple conditions or comorbidities in the same patient, and developing models for mild disease status as well as gathering evidence for diseases not included in the model, such as ones that may occur in future pandemics.

## Acknowledgments

We thank Akshay Balsubramani and members of the Kundaje and Boyd labs for helpful discussions. We also thank Stanford Health Care Clinical Virology Laboratory members Fumiko Yamamoto, Malaya K. Sahoo, ChunHong Huang, and Daniel Solis. We also thank the Stanford Covid-19 Biobank Study Group’s members: Elizabeth J. Zudock, Marjan M. Hashemi, Kristel C. Tjandra, Jennifer A. Newberry, James V. Quinn, Rosen Mann, Anita Visweswaran, Thanmayi Ranganath, Jonasel Roque, Monali Manohar, Hena Naz Din, Komal Kumar, Kathryn Jee, Brigit Noon, Jill Anderson, Bethany Fay, Donald Schreiber, Nancy Zhao, Rosemary Vergara, Julia McKechnie, Aaron Wilk, Lauren de la Parte, Kathleen Whittle Dantzler, Maureen Ty, Nimish Kathale, Arjun Rustagi, Giovanny Martinez-Colon, Geoff Ivison, Ruoxi Pi, Maddie Lee, Rachel Brewer, Taylor Hollis, Andrea Baird, Michele Ugur, Drina Bogusch, Georgie Nahass, Kazim Haider, Kim Quyen Thi Tran, Laura Simpson, Michal Tal, Iris Chang, Evan Do, Andrea Fernandes, Allie Lee, Neera Ahuja, Theo Snow, James Krempski.

## Author contributions

M.E.Z., A.K., and S.D.B. conceived the study and wrote the manuscript with input from all authors. *Data generation:* N.R., T.D.P., E.S.P., S.R.M., W.D., E.M.D., C.C.R., M.A.M., B.F.H., J.D.G., J.R.H., K.C.N., B.A.P., C.A.B., S.E.H., K.J., E.M., I.B., P.J.U., J.T.M., J.M.G., J.A.J., and S.Y. provided blood samples, as well as clinical and demographic data annotation and analysis and helpful discussions. J.Y.L., K.D.N., R.A.H., K.R., and B.L. prepared and sequenced samples. K.M.R. designed and created the data warehouse. M.E.Z. and K.R. ran the bioinformatics pipeline. M.E.Z. processed the external cohorts. *Model development*: M.E.Z. and S.D.B. developed Models 1 and 2. E.C., M.E.Z., J.K.M., S.D.B., R.T., and A.K. designed the Model 3 sequence and patient classification stages. M.E.Z. and J.K.M. evaluated language model embeddings for Model 3. M.E.Z., S.D.B., R.T., and A.K. designed the ensemble model. M.E.Z., J.K.M., and N.S. wrote the Python package. *Computational analysis:* M.E.Z. evaluated performance by dataset, disease, gene locus, and model component. N.S. compared SLE predictions and SLEDAI scores. J.K.M. and M.E.Z. performed the healthy donor replicate analysis. M.E.Z., R.T., and A.K. evaluated demographic influences on immune repertoires. M.E.Z., E.C., and A.K. analyzed the Shapley feature importances. M.E.Z., N.S., E.C., and R.T. analyzed the known binders.

## Data and materials availability

Prior published datasets are listed in **table S1**. New datasets will be deposited at publication time. The use of data was approved by Stanford University IRBs #13952, #48973, and #55689, as well as institutional approvals at local sites. Code is deposited online at https://github.com/maximz/malid.

## Materials and Methods

### B and T cell repertoire sequencing

We assembled immune receptor repertoires from 63 Covid-19, 95 chronic HIV-1, 86 Systemic Lupus Erythematosus (SLE), and 92 Type-1 Diabetes (T1D) patients, along with 217 healthy controls and 37 influenza vaccination recipients. Most non-Covid-19 cohort samples were collected before the emergence of SARS-CoV-2, except for the influenza vaccine cohort and some of the diabetes cohort and associated healthy controls. Covid-19 samples were collected early in the pandemic. Among Covid-19 patients, we excluded mild cases, samples prior to seroconversion, and patients known to be immunosuppressed. These filters limited model training data to active disease samples to improve our chances of learning patterns for the disease-specific minority of receptor sequences. However, we wanted to avoid creating an artificially simple classification problem from filtering to trivially separable immune states. To this end, we included both treatment-naive and treated SLE patients, and our HIV cohort included patients regardless of whether they generated broadly neutralizing antibodies to HIV. Had we instead restricted our analysis to HIV-infected individuals who produce broadly neutralizing antibodies, we may have created a more easily separable HIV class, due to the unusual characteristics of those antibodies (*18*).

Across these diverse immune states, over 16.2 million B and 23.5 million T cell receptor clones were sampled, PCR amplified with immunoglobulin and T cell receptor gene primers, and sequenced as previously described (*18*, *59*). Briefly, we amplified T cell receptor beta chains and each immunoglobulin heavy chain isotype in separate PCR reactions using random hexamer-primed cDNA templates, and performed paired-end Illumina MiSeq sequencing. To reduce the potential for batch effects, data collection followed a consistent protocol. Only IgH sequencing was performed for some older cohorts processed before the study was extended to include TRB sequencing. We annotated V, D, and J gene segments with IgBLAST v1.3.0, keeping productive rearrangements only (*86*). Using IgBLAST’s identification of mutated nucleotides, we calculated the fraction of the IGHV gene segment that was mutated in any particular sequence; this is the somatic hypermutation rate (SHM) of a B cell receptor heavy chain. On the other hand, T cell receptors are known not to exhibit somatic hypermutation. We also restricted our dataset to CDR-H3 and CDR3β segments with eight or more amino acids; otherwise the edit distance clustering method below might group short but unrelated sequences.

We grouped nearly identical sequences within the same person into clones, as described previously (*18*). To do so, for each individual, we grouped all nucleotide sequences from all samples (including samples at different timepoints) across all isotypes, and ran single-linkage hierarchical clustering to infer clonal lineages, iteratively merging sequence clusters from the same individual with matching IGHV/TRBV genes, IGHJ/TRBJ genes, and CDR-H3/CDR3β lengths, and with any cross-cluster pairs having at least 95% CDR3β sequence identity by string substitution distance, or at least 90% CDR-H3 identity, which allows for BCR somatic hypermutation (*18*).

We used the clonal lineage groupings to deduplicate the dataset. For each replicate of a sample from a patient, we kept one copy of each clone per isotype — choosing the sequence with the highest number of RNA reads. Similarly, we kept one copy of each TCRβ clone. Any replicates with fewer than 100 IgG, 100 IgA, and 500 IgD or IgM clones, or with fewer than 500 TRB clones, were rejected.

Among BCR sequences, we kept only class-switched IgG or IgA isotype sequences, and non-class-switched but still antigen-experienced IgD or IgM sequences with at least 1% SHM. By restricting the IgD and IgM isotypes to somatically hypermutated BCRs only, we ignored any unmutated cells that had not been stimulated by an antigen and were irrelevant for disease classification. The selected non-naive IgD and IgM receptor sequences were combined into an IgM/D group.

On average, any two patients had 0.0003% IgH and 0.166% TRB sequence overlap, underscoring the enormous diversity of T cell receptor and especially B cell receptor sequences, as would be expected from random sequence generation by the V(D)J recombination process followed by additional BCR somatic hypermutation.

### Cross-validation

We divided individuals into three stratified cross-validation folds, each split into a training set and a test set (**fig. S2**). Each individual was assigned to one test set. Some patients had multiple samples; all were grouped together for the cross-validation divisions. The splits were respected across the training of the complete *Mal-ID* pipeline. Stratified cross-validation preserved the global imbalanced disease class distribution in each fold. We also carved out a validation set from each training set. What remained of the training set was further subdivided into two parts we call “train-1” and “train-2”. The repertoire classification, CDR3 clustering, and language model base classifiers were trained on the training set and evaluated on the validation set. Then using the base models with highest validation set performance, the ensemble model was trained on the validation set, and then evaluated on the test set. In the case of multi-stage models like Models 2 and 3, the sequence classification stage was fit on the train-1 set, then the patient level aggregation stage was fit on train-2. When we used logistic regression classification models, regularization hyperparameters were tuned with additional nested cross-validation. This training process happens separately for each fold; in other words, one collection of models is trained using fold 1’s training, validation, and test sets, then a separate set of models is trained using fold 2’s training, validation, and test sets, and so on. On average in any fold, we observed 0.05% of IgH and 5.3% of TRB sequences shared between any pair of the train, validation, and test sets.

Since any single repertoire contains many clonally related sequences, but is very distinct from other people’s immune receptors, we made sure to place all sequences from an individual person into only the training, validation, or the test set, rather than dividing a patient’s sequences across the three groups. Otherwise, the prediction strategies evaluated here could appear to perform better than they actually would on brand-new patients. Given the chance to see part of someone’s repertoire in the training procedure, a prediction strategy would have an easier time of scoring other sequences from the same person in a held-out set. Had we not avoided this pitfall, models may also have been overfitted to the particularities of training patients. For the minority of individuals with multiple samples, we accordingly made sure that, in each cross-validation fold, all samples from the same person were grouped together into one of the training, validation, or test sets, as opposed to being spread across multiple sets. This principle was also respected for all nested cross-validation.

Finally, for the purpose of external cohort validation, we repeated the model training procedures with a “global” fold designed to incorporate all the data, by having only a training set and a validation set but no test set (**fig. S2**). Repertoires from independent external studies are used in place of the test set at evaluation time.

### Evaluation metrics

Models were trained with the python-glmnet implementation of logistic regression (with multinomial loss and regularization strength tuned through cross-validation), as well as with the scikit-learn implementations of random forests (with 100 trees) and support vector machines (in “each class versus the rest” mode, with linear kernel and default regularization strength hyperparameter C=1.0). In all cases, we used prevalence-balanced class weights inversely proportional to input class frequencies. Predicted labels from all test sets were concatenated for global accuracy evaluation. Performance metrics that take predicted class probabilities as input, including AUROC and AUPRC, were computed separately for each fold, because probabilities may be on different scales in each fold and should not be combined into a global AUROC or AUPRC score. For overall performance, we report multi-class AUROC and AUPRC calculated in a one-versus-one fashion, taking the class size-weighted average of the binary AUROCs/AUPRCs calculated for each pair of classes, allowing each class a turn to be the positive class in the pair. For each disease class’s individual performance, we report multi-class AUROC calculated in a one-versus-rest fashion. The AUROC and AUPRC measures do not reflect classification abstention, because abstained samples have no predicted class probabilities and cannot be included in the computation of metrics that use predicted probabilities. On the other hand, every abstention hurts label-based metrics like accuracy: each abstention counts as a prediction error. All analyses were performed and plotted with software versions *python v3.9.17, numpy v1.24.3, pandas v1.5.3, scipy v1.11.1, scikit-learn v1.2.2, python-glmnet v2.2.1, pytorch v2.0.1, bio-transformers v0.1.17, matplotlib v3.7.1*, and *seaborn v0.12.2*.

### Architecture overview

#### Model 1: Overall repertoire composition

The first machine learning model uses an individual’s IgH or TRB repertoire composition to predict disease status. Prior studies have reported immune status classification using deviations in B cell or T cell V(D)J recombination gene segment usage from healthy individuals (*23*, *87*, *88*). Certain V gene segments may be more prevalent among antigen-responding V(D)J rearrangements than in the population of immune receptors in naïve lymphocytes, and these gene segments increase in frequency as antigen-specific cells become clonally expanded (*70*, *89*), which can be seen in our data (**fig. S7a**). We previously identified class-switched IgH sequences with low somatic mutation (SHM) frequencies as prominent features of acute infection with Ebola virus or SARS-CoV-2, consistent with naïve B cells recently having class-switched during the primary response to infection (*70*, *89*, *90*). V gene usage changes and other repertoire changes have also been described in chronic infectious or immunological conditions (*14*, *18*). Therefore, we trained a logistic regression model with V/J gene counts, along with somatic hypermutation rate for IgH data, as features.

#### Model 2: Convergent clustering of antigen-specific sequences by edit distance

The second classifier detects highly similar CDR3 amino acid sequences shared between individuals with the same diagnosis, an approach we and others have previously reported (*17*, *18*, *21*, *22*). The CDR3s are the highly variable regions of IgH and TRB that often determine antigen binding specificity. For each locus, we clustered CDR3 sequences with the same V gene, J gene, and CDR3 length that had high sequence identity, allowing for some variability created by somatic hypermutation in B cell receptors. A new sample’s sequences can then be assigned to nearby clusters with the same constraints. We selected clusters enriched for sequences from subjects with a particular disease, using Fisher’s exact test and setting a significance threshold based on cross-validation with data derived from different individuals. These clusters represent candidate sequences predictive of a specific disease across individuals. To score a new sample, we assigned its sequences to the identified predictive clusters. For each sample, we counted how many clusters associated with each disease were matched, and used these counts as features in a logistic regression model to predict immune status.

#### Model 3: Immune receptor sequence features extracted from a large language model

Small changes to immune receptor amino acid sequences can alter receptor structure and function, while different structures with divergent primary amino acid sequences can bind the same target epitope (*91*, *92*). We used a protein language model, which transforms BCR and TCR amino acid sequences into a lower-dimensional representation, to estimate functional similarities between sequences that extend beyond sequence alignment. Specifically, we used ESM-2, a self-supervised model trained to predict masked amino acids from the remaining sequence context of a protein, learning complex statistical relationships between residues in each sequence and encoding functional and evolutionary relationships across sequences (*45*, *93*). Prior autoencoder models, which also convert immune receptor sequences to a latent representation, have enabled classification and clustering of functionally related sequences (*39*, *41*). However, ESM-2 is a large language model with significantly more parameters that is trained on a much larger compendium of over 65 million proteins across the tree of life, which allows it to learn richer latent representations that encode properties of a broad diversity of protein structures and functions (*45*, *93*). We developed machine learning models with a novel two-stage training strategy to predict patient-level disease status based on ESM-2-derived representations of their immune repertoire. First, we trained machine learning models to map ESM-2 derived 640-dimensional latent representations of each receptor sequence from each patient sample to a surrogate disease state corresponding to the disease state of the patient. Each model is specialized to one IGHV gene and isotype combination in the BCR case, or to one TRBV gene in the TCR case. Somatic hypermutation rate was used as an additional feature in the BCR case (hypermutation does not occur in TCRs). Then we trained a second-stage model that aggregates predicted probabilities of disease state of all sequences in a patient sample, again grouped by IGHV gene and isotype or by TRBV gene, to predict disease state at the patient level.

#### Ensemble of B and T cell models

Finally, we combined all three classifiers (overall repertoire composition, clustering by edit distance, and language model representation) for IgH and three for TRB into the final *Mal-ID* ensemble predictor of disease (**fig. S1**). As with the individual component models in *Mal-ID*, we trained a separate metamodel for each cross-validation group, maintaining strict separation of each individual’s data into training, validation or test datasets.

### Model 1: Disease classifier using overall BCR or TCR repertoire composition features

For each sample, we created IgG, IgA, IgM/D, and TRB summary feature vectors by tallying IGHV/TRBV gene and IGHJ/TRBJ gene usage, counting each clone once. We ranked IGHV or TRBV genes by training set prevalence and excluded the bottom half, to avoid overfitting to minute differences in rare V gene proportions between cohorts. To account for different total clone counts across samples, we normalized total counts to sum to one per sample. Then we log-transformed and Z-scored (i.e. subtracted the mean and divided by the standard deviation, to achieve zero mean and unit variance) the matrix representing how counts are distributed across V-J gene pairs. Finally, we performed a PCA to reduce the count matrix to fifteen dimensions. All transformations were computed on each training set and applied to the corresponding validation and test sets. In addition, for each sample’s subset of BCR sequences belonging to each isotype, we calculated the median sequence somatic hypermutation rate and the proportion of sequences that are somatically hypermutated (with at least 1% SHM). Only BCRs have somatic hypermutation, so we did not include mutation rate features of TCRs. In total, we arrived at 51 features across IgG, IgA, and IgM/D (fifteen count matrix principal components and two mutation rate features per isotype) for the IgH repertoire composition model, and 15 features for the TRB repertoire composition model.

We fit separate logistic regression linear models with L1 regularization on the 51-dimensional (17 x 3 isotypes) BCR and 15-dimensional TCR feature vectors from each sample to predict disease. Features were standardized to zero mean and unit variance. We repeated this feature engineering and model training procedure on each cross-validation fold separately. The best performing models, according to average validation set AUROC across three cross-validation folds for the disease classification task on our primary dataset, were elastic net logistic regression with an L1/L2 regularization ratio of 0.25 for BCR and lasso logistic regression for TCR.

### Model 2: Disease classifier by clustering CDR-H3 sequences with edit distance

We performed single-linkage clustering on CDR3β sequences from T cells with identical TRBV genes, TRBJ genes, and CDR3β lengths, and separately on CDR-H3 sequences from B cells with identical IGHV genes, IGHJ genes, and CDR-H3 lengths, as described previously (*18*). Nearest-neighbor clusters were iteratively merged if any cross-cluster pairs had high sequence identity: at least 90% for CDR3β or 85% for CDR-H3, allowing for somatic hypermutation in B cells, as measured by string substitution distance (normalized Hamming distance). Clustering was performed on the train-1 data sets. This process was run separately for each cross-validation fold.

#### Filter to BCR and TCR disease-specific enriched clusters

For each sequence cluster found in the train-1 portion of a cross-validation fold’s training set, we performed a Fisher’s exact test using a two-by-two contingency table denoting how many unique people have a particular disease and have some receptor sequences fall into the cluster. In other words, each cluster’s p value from the Fisher’s exact test denotes the cluster’s enrichment for a particular disease. This approach is consistent with prior work that selects a set of disease-specific enriched sequences, then counts exact matches to this sequence set in new samples (*17*). Given a p value threshold, the full list of training set clusters was filtered to clusters specific for each disease type. We performed all the following featurization and model fitting steps for p values ranging from 0.0005 to 0.05, then selected the p value that led to the highest train-2 set performance as measured by the Matthews correlation coefficient (MCC) score, a classification performance metric that is well-suited to imbalanced datasets (*94*). The final chosen p values differed depending on the cross-validation fold and the receptor type (i.e. BCR or TCR).

#### Compute BCR and TCR cluster membership feature vectors for each sample

For each selected enriched cluster, we created a cluster centroid: a single consensus sequence. Recall that each cluster member is a clone from which only the most abundant sequence was sampled. Rather than having each cluster member contribute equally to the consensus centroid sequence, contributions at each position were weighted by clone size, the number of unique BCR or TCR sequences originally part of each clone. Sequences from a sample were then matched to these predictive cluster centroids. In order to be assigned, a sequence must have the same IGHV/TRBV gene, IGHJ/TRBJ gene, and CDR-H3/CDR3β length as the candidate cluster, and must have at least 85% (BCR) or 90% (TCR) sequence identity with the consensus sequence representing the cluster’s centroid. After assigning sequences to clusters, we counted cluster memberships across all sequences from each sample. Cluster membership counts were arranged as a feature vector for each sample: a sample’s count for a particular disease was defined as the number of disease-enriched clusters into which some sequences from the sample were matched. This featurization captures the presence or absence of convergent T cell receptor or immunoglobulin sequences (separated by locus, but without regard for IgH isotypes).

#### Fit and evaluate model for each locus

Features were standardized, then used to fit separate BCR and TCR logistic regression models mapping from cluster counts to patient diagnosis. The models were fit on each train-2 set and evaluated on the corresponding validation set. The best performing models, according to average validation set AUROC across three cross-validation folds for the disease classification task on our primary dataset, were ridge logistic regression for BCR and lasso logistic regression for TCR.

We abstained from prediction if a sample had no sequences fall into a predictive cluster; this indicated no evidence was found for any particular class. Abstentions hurt accuracy and MCC scores, but were not included in the AUROC calculation, since no predicted class probabilities are available for abstained samples. Fewer than 3% of samples resulted in abstention (**table S2**).

#### Comparison to exact matches approach

Briefly, Emerson et al. classified cytomegalovirus (CMV) exposure by counting the number of TRB sequences that were exact matches to a CMV-associated list derived from a training set of CMV+ and CMV-individuals (*17*). CMV-associated sequences were determined with a Fisher’s exact test using a two-by-two contingency table denoting how many unique people are CMV+ and have a particular sequence; the threshold on Fisher’s exact test p values was selected by cross-validation.

We re-implemented this method for the *Mal-ID* dataset to compare the “exact sequence matches” featurization of Emerson et al. against the “fuzzy matches” featurization of the CDR3 clustering component of *Mal-ID*. The binary classification generative model used in Emerson et al. after the featurization step does not translate to our multi-class disease classification problem, so we instead used the same classification framework as the CDR3 clustering model: each sample’s feature vector consisted of the number of disease-specific hits for each disease, normalized by the total size of the sample. Additionally, we ensured that both models had a consistent approach to abstention. The CDR3 clustering model abstains on samples that had zero matches to any disease-associated cluster; similarly, our implementation of Emerson et al. in the multi-class problem abstains on samples that had zero matches to any disease-associated sequence (i.e. there is no evidence of disease). Just as when training the CDR3 clustering model, the exact matches featurization and model fits were performed for different p value thresholds, then the best threshold was chosen by optimizing performance on the second part of the training set (train-2) using the MCC score. Therefore, the Emerson et al. and CDR3 clustering models are trained the same way in this comparison, differing only in whether the featurization step finds exact sequence matches or fuzzy matches.

### Model 3: Disease classifier using language model embeddings

The analysis pipeline for classifying disease with language model embeddings of sequences is complex, but necessarily so because it aggregates individual sequence data to generate patient-level predictions.

#### Generate embeddings

We embedded the CDR-H3/CDR3β segments of each receptor sequence with the 30-layer, 150-million-parameter ESM-2 neural network (*45*), using the bio-transformers v0.1.17 implementation. A final 640-dimensional vector representation was calculated by averaging ESM-2’s hidden state over the original protein’s length dimension.

#### Train sequence-level disease classifier for each sequence category

First, we trained classification models to map sequences to disease labels — one model per fold and per sequence category, defined as an IGHV gene and isotype pair for BCR sequences or a TRBV gene for TCR sequences. As input data, we used ESM-2 embeddings (standardized to zero mean and unit variance), along with somatic hypermutation rate in the BCR case. To train the individual-sequence-level model, we provided noisy sequence labels derived from patient global immune status. This is because the available ground truth data associates *patients*, not sequences, with disease states; we do not know which of their sequences are truly disease related. Since we have no true sequence labels, we also cannot evaluate classification performance for the sequence-level classifier directly. These sequence-level classifiers were trained on the train-1 set of each cross-validation fold.

#### Aggregate sequence predictions within each sequence category

We combined predictions for individual BCR or TCR sequences into a patient sample-level prediction by the following procedure. Given a sample with *n* BCR (or TCR) sequences, we first scored each sequence with the corresponding sequence model. For example, we applied the IGHV3-53, IgG model to input sequences arising from the IGHV3-53 gene segment and the IgG isotype. Each sequence now has a vector of *k* predicted probabilities, with one value for each of the *k* disease classes. These values are only comparable between sequences that were scored by the same model, as models for different sequence groups are not guaranteed to have matching calibration. Therefore, we next aggregated predicted class probabilities among sequences from the same sequence category, one IGHV gene and isotype (or one TRBV gene) at a time. To calculate the aggregate probability for each of the *k* classes, we used one of the following methods:

- Mean
- Median
- Trimmed mean: Remove the lowest 10% of sequence-level probabilities before calculating the mean.
- Entropy thresholded mean: Before taking the mean, remove any sequences whose predicted class probability vectors had high entropy, indicating they carry little information that could indicate a particular disease class. A sequence with probabilities of *1/k* for all *k* classes would have the highest possible entropy. We removed sequences whose entropy was within either 10% or 20% of this maximal value.

This procedure gives the final *k*-dimensional predicted disease class probabilities vector for each sequence category in each sample. For example, it computes P(Covid19) among IGHV1-24/IgG sequences, P(HIV) among IGHV1-24/IgG sequences, and so on; then similarly P(Covid19) among IGHV3-53/IgA sequences, P(HIV) among IGHV3-53/IgA sequences, and so forth.

#### Map from aggregate predictions for each sequence category to a sample prediction

Using the aggregated sequence-level predictions, we make a final prediction for the sample with a second-stage model. This model was fitted in a one-versus-rest fashion, and the submodel for each class was trained only with features corresponding to that class. For example, the Covid-19-vs-rest model was provided P(Covid-19) in IGHV1-24/IgG, P(Covid-19) in IGHV3-53/IgG, and so on, but not P(HIV), P(Influenza), P(Lupus), P(T1D), or P(Healthy). This design prohibits unwanted feature leakage: deciding whether a sample is from a Covid-19 patient should rely only on sequence-level probabilities for the Covid-19 class, not any other classes. Also, we incorporated features for only the top 50% of IGHV or TRBV genes to avoid having far more features than samples for this second-stage model, and because rare V genes may not be present in all samples. Therefore, the number of features in this second-stage model for the BCR case was half the number of IGHV genes, times three isotype categories: IgG, IgA, and IgM/D excluding naive B cells with <1% somatic hypermutation. For TCR, which has no isotype subdivisions, the number of features was half the number of TRBV genes. Each sample’s features were reweighed according to sequence category frequencies. In the BCR case, frequencies were computed separately for each isotype to account for technical variation in isotype frequencies between sequencing runs. The aggregation model was trained on the train-2 set in each cross-validation fold.

#### Evaluate classifier

We evaluated the pipeline by computing sample-level classification performance on the validation set using AUROC scores. (The one-versus-rest model predicted probabilities are not necessarily calibrated against each other, so we did not evaluate accuracy or other metrics determined by the comparison of predicted class probabilities for selecting a winning label). For the BCR case, the highest validation set performance on our primary dataset was achieved by a pipeline consisting of random forest sequence-level models, followed by a random forest second-stage model using mean aggregation. In the TCR case, the best pipeline used one-versus-rest ridge logistic regression sequence-level models, with a random forest second-stage model using mean aggregation after an entropy cutoff at 20% below the maximal entropy value (**table S7**). To evaluate feature contributions to predictions of each disease class, we ran Tree SHAP on each one-class-versus-rest random forest aggregation model, and averaged the SHAP feature importance values across positive class instances from the train-2 data used to train the aggregation model. SHAP values were rescaled from 0 to 1. Alternatively, to find SHAP clusters, we performed Louvain clustering (resolution 1.0) on the full SHAP value matrix in which rows represent positive class examples and columns represent features, then calculated average SHAP values within each cluster.

### Ensemble metamodel

After training repertoire composition, CDR3 clustering, and language model embedding models on each fold’s training set, we combined the classifiers with an ensemble strategy. For each fold, we ran all trained base classifiers on the validation set, and concatenated the resulting predicted class probability vectors from each base model. We carried over any sample abstentions from the CDR3 clustering model (the other models do not abstain). Finally, we trained a ridge logistic regression classification metamodel to map the combined predicted probability vectors to validation set sample disease labels. We evaluated this metamodel on the held-out test set. To evaluate individual model component contributions, we refit the metamodel with subsets of features, such as only those features derived from models 1 and 2.

### Batch effect evaluation using language model embeddings

Having integrated many datasets in this study, we sought to test whether our disease classification performance was driven by technical differences between batches of library preparation or sequencing instrument run. It would be expected in any study of human cohorts to identify some batch effects, given the difficulty of collecting identical samples in identical manner, at identical severity and timepoints, from patients suffering from diseases that appear in different populations at different frequencies. Notably, the IgH data collected for individual participants in this study were typically based on multiple Illumina MiSeq sequencer runs, and were combined prior to analysis. Many of our sequencing run batches included only one disease type, but batches that included both diseased and healthy controls from the same population permitted accurate classification of the disease or healthy state, for example, with classification of HIV-infected patients and healthy controls that were sequenced together in the same batch, or SLE patients and healthy controls sequenced in the same batch.

Acknowledging that there were biological differences between many sequencing batches that were enriched for a particular disease state, and that several sequencer runs were performed for some sample sets, we evaluated the potential impact of these batch differences using the language model embeddings of BCR and TCR repertoires from the disease types found in multiple batches: Covid-19 patients, SLE patients, and healthy donors. We applied the kBET batch effect metric from the single cell sequencing literature (*95*, *96*). kBET measures whether cells from many batches are well-mixed by comparing the batch label distribution among each cell’s neighbors to the global distribution. In place of cells described by gene expression vectors, we have sequences described by language model embedding features. We measured kBET for every disease in every test set fold and in both BCR and TCR data. For example, we constructed a k-nearest neighbors graph (k = 50) with all BCR sequences from Covid-19 patients in test fold 1. We performed chi-squared tests for the difference between the batch label distribution among each sequence’s 50 nearest neighbors and the expected distribution from the total number of sequences belonging to each batch in the entire graph. After multiple hypothesis correction with a significance threshold of p=0.05, we measured the number of sequences for which we could reject the null hypothesis that the local neighborhood batch distribution is the same as the global batch distribution. Aggregating these results by disease across gene loci and folds, we see that the null hypothesis is rejected for only 18.2% of sequences on average, suggesting that the sequence data in the graph are well mixed according to batch (**table S8**). The average rejection rate is higher for Covid-19 BCR sequences at 44.1%, which may be influenced by disease severity differences between cohorts (**table S1**). Time point differences between batches may also influence kBET metrics for acute diseases like Covid-19. At earlier time points, Covid-19 patient repertoires may include more healthy background sequences, leading to a different batch overlap graph in comparison to how batches compare after clonal expansion of Covid-19 responding sequences. Overall, these results suggest that most sequences have well-mixed batch proportions amongst their nearest neighbors.

### Validation on external cohorts

The best test of whether our model has learned true biological signal as opposed to batch effects is whether our model generalizes to unseen data from other cohorts. For the purposes of evaluating external cohorts, rather than using models trained on our cross-validation divisions of the data, we trained a set of “global” models incorporating all *Mal-ID* data without holding out a test set (**fig. S2**). To train the ensemble metamodel, we still held out a validation set, with a ratio of training set to validation set size equivalent to the ratio used in the cross-validation regime.

We downloaded data from other BCR and TCR Covid-19 patient and healthy donor repertoire studies with cDNA sequencing (*54–58*, *97*). Among acute Covid-19 cases, we selected active disease timepoint samples at least two weeks after symptom onset, after which time we would expect seroconversion (*70*). We reprocessed sequences through the same version of IgBLAST and IgBLAST reference data used for the primary *Mal-ID* cohorts, to ensure consistent gene nomenclature. (This was not possible for the Britanova et al. datasets (*57*, *58*) because the raw sequences were unavailable, so we used their gene calls and confirmed the naming was consistent with our training data, especially for indistinguishable TRBV genes TRBV6-2/6-3 and TRBV12-3/12-4.) We embedded productive CDR3 sequences with the language model, then processed the downloaded repertoires through the entire *Mal-ID* model architecture. We also tuned class decision thresholds to adapt the model to the new base rates of disease in the data. Specifically, we held out several external cohort samples and reweighted their predicted class probabilities to optimize the MCC score. After this procedure, the winning label for each sample is chosen based on the class with highest predicted probability after class weights are applied. If a class had its probabilities reweighted by 1/5, for example, the model must be five times more confident to choose that class label. This procedure affected only the confusion matrix, accuracy, and other metrics based on predicted labels.

Additionally, we retrained *Mal-*ID after downloading TCR repertoire data collected with the Adaptive Biotechnologies genomic DNA sequencing protocol (**table S4)**. This data was reprocessed with the same IgBLAST version as above, for consistency.

### Predicting demographic information from healthy subject repertoires

We repeated the model training process to predict age, sex, or ancestry instead of disease. Input data was limited to healthy controls to avoid learning any disease-specific patterns. To cast this as a classification problem, age was discretized either into deciles, as a binary “under 50 years old” / “50 or older” variable, or as a binary “under 18 years old” / “18 or older” variable. Only one healthy control individual was over 80 years old, therefore our data do not assess repertoire changes at more extreme older ages. We excluded the healthy individual over 80 years old from the analysis.

For each of the demographic prediction tasks, we trained the full BCR+TCR *Mal-ID* architecture on all cross-validation folds. We note that we did not explicitly introduce data from allelic variant typing in germline IGHV, IGHD, or IGHJ gene segments or in HLA genes into our models, but such data could be expected to increase detection of ancestry in such datasets.

### Evaluating predictive power of potential demographic confounding variables

We retrained the entire *Mal-ID* disease-prediction set of models on the subset of individuals with known age, sex, and ancestry. (As above, we excluded any individuals over 80 years old.) Additionally, we regressed out those demographic variables from the feature matrix used as input to the ensemble step. Specifically, we fit a linear regression for each column of the feature matrix, to predict the column’s values from age, sex, and ancestry. The feature matrix column was then replaced by the fitted model’s residuals. This procedure orthogonalizes or decorrelates the metamodel’s feature matrix from age, sex, and ancestry effects. We regressed out covariates at the metamodel stage because it is a sample-level, not sequence-level model, and age/sex/ancestry demographic information is tied to samples rather than sequences.

Separately, we also trained models to predict disease from either age, sex, or ancestry information encoded as categorical dummy variables. Here, no sequence information was provided as input. Finally, we trained metamodels with both demographic features and sequence features, along with interaction terms between the demographic and sequence features to allow for interaction effects. Comparing the performance of these models to the demographics-only models shows the added value of adding sequence information.

### Model ranking of known antigen-specific sequences

We downloaded the June 13, 2023 version of CoV-AbDab (*75*), and reprocessed these B cell receptor heavy chain sequences through the same version of IgBLAST used for our primary cohorts to ensure consistent V gene nomenclature. However, CoV-AbDab contains amino acid sequences, rather than nucleotide sequences as in our internal data, so we used the protein version of IgBLAST (“igblastp”) and quantified somatic hypermutation based on the percentage of mutated amino acids. We filtered to antibody sequences known to bind to SARS-CoV-2 (including weak binders, but excluding sequences shown to selectively bind certain viral variants but not others), and only kept sequences from human patients or vaccinees. We clustered the selected SARS-CoV-2 binders with identical IGHV gene, IGHJ gene, and CDR-H3 lengths and at least 95% sequence identity, using single linkage clustering as in the pipeline for our primary cohorts. As a result, several related sequences were combined and replaced by a consensus sequence. This preprocessing was repeated for influenza-specific antibody sequences from human patients and vaccinees (*78*), excluding H5N1 and H7N9 vaccine or infection data because those strains are not included in the seasonal flu vaccine that our classifier was trained to distinguish.

Similarly, we downloaded the ImmuneCode MIRA database (*79*), version 002.1, and reprocessed these T cell receptor beta chain sequences with our pipeline’s standard IgBLAST version for consistent V gene nomenclature. As above, we filtered to productive sequences from patients with acute Covid-19, and also to only the TRBV genes present in our dataset, as any others would not be compatible with the sequence model, which uses V gene segment identity as a feature. Among the remaining SARS-CoV-2 associated sequences, we deduplicated those with identical TRBV genes, TRBJ genes, and CDR3β sequences.

We scored the external databases of known binder sequences using Models 2 and 3 trained on the global fold. Isotype designations were not available in the BCR antigen-specific datasets; we applied our IgG sequence models because many antigen-specific B cells in Covid-19 have been reported to express IgG (*70–73*). Correspondingly, we compared to IgG sequences from healthy donors in the global fold’s validation set, which were held out from training. To perform the statistical test shown for a particular V gene (e.g. IGHV1-24 for the Covid-19 analysis), we conducted a one-sided permutation test to assess whether known binder sequences had higher model 3 predicted Covid-19 class probabilities compared to sequences from healthy individuals. The permutation test ensured that all sequences originating from each healthy donor individual retained their grouping (i.e. had consistent binder/non-binder labels) throughout the process of performing 1000 label permutations. Since the known binders have low prevalence and since permutation affects the prevalence, we computed the AUPRC fold change over baseline prevalence in each permutation, then calculated the p-value as the proportion of permutations whose AUPRC fold change was greater than the observed AUPRC fold change in the original data.

## Supplementary Text

### Sequence category contributions to disease predictions

In discriminating between different diseases, sequences prioritized for Covid-19 prediction used IGHV gene segments seen in independently isolated antibodies that bind SARS-CoV-2 spike antigen: IGHV1-2 (*98–102*), IGHV1-46 (*103*, *104*), IGHV2-70 (*105*), IGHV3-7 (*106*, *107*), IGHV3-11 (*99*), IGHV3-30 (*73*, *101*, *108–111*), IGHV3-53 (*112*), IGHV4-4 (*113*), IGHV4-31 (*114*), and IGHV4-39 (*109*) (**Fig. 3A**). Similarly, IGHV1-24, found in a prominent class of N-terminal domain-directed antibodies (*115*, *116*), was highly ranked, as was IGHV4-59 from nucleocapsid binding antibodies (*109*). A gene segment known to produce autoreactive antibodies, IGHV4-34 (*117*), also had high classification priority, matching earlier reports of higher prevalence in Covid-19 patients (*118*).

IGHV4-34 was also prioritized for SLE prediction along with IGHV4-59 (**Fig. 3D**), matching prior reports of higher frequency expression of these gene segments in SLE patients (*14*, *119*); they have also been reported as the source for several known autoantibodies (*120*). Additionally, the SLE model prioritizes IGHV1-69, IGHV3-7, and IGHV3-30, which have been found in anti-dsDNA antibodies specific to lupus (*121*, *122*). IGHV6-1, a gene known to encode anti-DNA and anti-phospholipid antibodies characteristic of lupus (*123*), also earned high priority. Some other V genes identified by the model, specifically IGHV1-18, IGHV1-46, IGHV3-9, and IGHV3-74, have been reported to have higher frequencies in lupus patients relative to rheumatoid arthritis patients (*124*), but others have not yet been characterized in the lupus literature to our knowledge: IGHV1-24, IGHV3-13, IGHV4-31, and IGHV4-61.

Similarly, the IGHV genes prioritized for T1D prediction have also not been associated with this disease in the literature to our knowledge: IGHV1-3, IGHV1-69, IGHV3-7, IGHV3-23, IGHV3-49, IGHV3-66, and IGHV3-74. IGHV genes prioritized in influenza class prediction, including IGHV1-18 (*125*, *126*), IGHV2-70 (*127*), IGHV3-7 (*128–130*), IGHV3-23 (*126*, *128*, *131–135*), IGHV3-30 (*128*), IGHV3-48 (*128*), IGHV3-66 (*136*), IGHV4-39 (*129*), IGHV4-59 (*20*, *128*), and IGHV5-51 (*130*, *137*), have been found in antibodies reactive to influenza virus. In HIV, IGHV4-34 has been described in HIV-specific B cell responses with unusually high somatic hypermutation frequencies in individuals producing broadly-neutralizing antibodies (*18*). It was ranked highly for HIV classification by the model (**Fig. 3B**). So were IGHV1-2, the source of VRC01-class broadly-neutralizing antibodies (*138*, *139*), and IGHV3-30, which has been found in other broadly-neutralizing antibodies (*140*). These V gene segments were not used far more commonly in the IgH germline loci of African populations, suggesting they are not prioritized due to the impact of demographic factors on immune repertoire data. This check is important because the HIV cohort was our largest African cohort and other genes, such as IGHV4-38-2, have different frequencies in this population (**fig. S15A**).

We repeated the analysis for TRBV gene contributions. As expected from genetic variation in the alleles of HLA proteins that restrict TCR binding, some TRBV genes were also stratified by ancestry (**fig. S15B**). TRBV10-2, TRBV24-1, and TRBV25-1, all gene segments enriched in African healthy controls, were among the most highly ranked TRBV gene groups for classifying our predominantly African HIV cohort (**fig. S16B**). However, TRBV5-1, TRBV6-1, TRBV7-2, and TRBV30 were also prioritized for HIV classification but were not enriched in African healthy controls. TRBV2, TRBV6-6, TRBV12-3, and TRBV18 were prioritized for T1D prediction (**fig. S16E**) and previously reported among islet antigen reactive T cells (*141*, *142*). However, TRBV12-3 and TRBV18 also had potentially age-associated differences in strength of contribution, which was seen when we decomposed the T1D Shapley feature importances into two clusters, one that is 71% composed of pediatric patients, and a second that is half pediatric and half adult (**fig. S13, C** and **D**). The clusters have distinct V gene prioritizations, indicating that different sequence signals identified the patients as positive for T1D. The lupus TRBV gene contributions can also be divided into two clusters, one of which is predominantly (88%) adult, while the other consists entirely of pediatric patients (**fig. S12, C** and **D**). Different TRBV genes are prioritized in the two clusters, and TRBV25-1 signal skews predictions to be more positive in the predominantly adult cluster but discourages lupus prediction in the pediatric cluster. These subtle differences suggest complex disease subtypes that may be divided by age or by treatment status, as the pediatric cohort is treatment naive. However, we did not see lupus or T1D cluster associations with age in the BCR model (**fig. S12, A** and **B**, **fig. S13, A** and **B**).

### Known binder sequence detection

Evaluating the sequence models specialized by IGHV gene, Model 3 AUROCs averaged 0.60 +/− 0.07, ranging up to 0.78, meaning known binders with certain IGHV usage were often ranked higher than healthy sequences by Covid-19 predicted probability (**Fig. 4, E** and **G**), despite no knowledge of these binding relationships during training. Since known binders have 2.7% prevalence when paired with healthy donor sequences, we also evaluated AUPRC, an alternative score for distinguishing the sequence classes within each IGHV gene that may be better suited to class-imbalanced settings (*143*, *144*). AUPRCs averaged 1.88-fold +/− 1.03 change higher than baseline prevalence, ranging up to 6.9-fold over baseline (**Fig. 4F**). IGHV1-24, IGHV2-70, and IGHV3-7 had high AUROCs and normalized AUPRCs, are reported in the Covid-19 literature, and also had high SHAP feature importance scores as described above (**Fig. 3**).

We also tested whether Model 2 identifies SARS-CoV-2 associated patterns by comparing Hamming distances from known binder and healthy donor sequences to Model 2’s Covid-19 associated clusters. During training, possible clusters are identified by clonal lineage parameters (IGHV gene, IGHJ gene, and CDR3 length), divided by Hamming distance, and kept only if highly enriched for sequences from patients with a particular immune state (determined by a statistical test with a strict p value threshold). This process leads to a small set of Covid-19 associated clusters. Therefore only 21% of individual binder sequences had clonal lineage parameters matching any Model 2 Covid-19 clusters, even though 97% of *patients* had sequences match Model 2’s clusters (**table S2**). Model 2 missing most known binders reflects its focus on finding shared public clones, whereas binding sequences may be private to an individual. Therefore, we evaluated Model 2’s ability to identify potential Covid-19 binders, not how well it rules them out. We called positives if query sequences matched any Covid-19 cluster’s clonal lineage parameters, evaluating each IGHV gene individually (consistent with Model 2’s division of sequences before clustering). Moderately high recall in some IGHV genes (average 15.5% +/− 15.0%, ranging up to 62.8%) suggests Model 2 usually does not overlook true binders, but low precision (average 4.2% +/− 4.4% across IGHV genes, ranging up to 19.0%) indicates the model also classifies many non-binders as binders (**Fig. 4, A** and **B**). Compared at equivalent precision, Model 3 generally had higher recall than Model 2 (**Fig. 4I**), indicating Model 3 captures more true binders at the cost of including false positives. However, Model 2 precision rose to 100% for key IGHV genes if identifying binders by 85% sequence identity to any Covid-19 sequence cluster (**Fig. 4, C** and **D**), the threshold from the main Model 2 pipeline and within the range used by other research groups (*145*). While this model was even more conservative, when it did predict sequences to be binders, those predictions were more likely to be correct. Some genes with perfect precision were IGHV1-24, IGHV3-7, IGHV3-30, IGHV3-53, and IGHV4-39 — all previously identified in the literature and having high SHAP values in Model 3 (**Fig. 3**, **Supplementary Text**). On the other hand, recall was low (maximum of 3.5%) because most sequences received negative predictions, raising the number of false negatives. Recall cannot be directly compared between Models 2 and 3 under the stricter 85% sequence identity decision threshold, because Model 2’s precision of exactly 0% or 100% in each IGHV gene falls at the boundaries of Model 3’s precision-recall curve. Despite low recall, Model 2 appears to have also learned true Covid-19 patterns, based on high precision in important IGHV genes.

Model 3 can score sequences that Model 2 cannot evaluate. For the 79% of known binders whose clonal lineage parameters do not match public clone clusters, Model 3 AUROCs averaged 0.59 +/− 0.06 across IGHV genes, ranging up to 0.75 (**Fig. 4H**), with AUPRCs averaging 1.63-fold +/− 0.46 change over baseline prevalence, with maximum 2.80-fold change. In this manner, Models 2 and 3 are complementary: Model 2 can confidently identify a subset of sequences as very similar to convergent clusters found in Covid-19 training set patients, and Model 3 can evaluate the remaining sequences.

In order to further test the conclusion that Models 2 and 3 learned sequence patterns truly associated with disease, we also compared scores between influenza known binders and healthy donor sequences (*78*). Here, we used influenza predictions generated by Models 2 and 3 after they were trained on samples from seasonal flu vaccine recipients. The influenza known binder dataset was smaller (0.62% prevalence when combined with healthy sequences). Model 3 AUROCs were 0.55 +/− 0.07 across IGHV genes, ranging up to 0.65 (**fig. S17, E** and **G**). Normalized AUPRCs were 1.95-fold +/− 1.00 change over baseline prevalence, up to a maximum of 4.00-fold change (**fig. S17F**). The AUROC and AUPRC scores were high for IGHV3-30, IGHV3-48, and IGHV4-39, all of which were described earlier because they have been reported in the influenza literature and earned high SHAP values in the Model 3 aggregation piece. On the other hand, Model 2, when calling positives based on having shared clonal lineage parameters with any influenza cluster, achieved low precision (maximum of 2.6%) and moderate recall (30.8% on average, ranging up to 51.5%; **fig. S17, A** and **B**). Comparing the models, recall was not consistently higher for Model 2 or Model 3 when evaluated at equivalent precision (**fig. S17I)**. Under the stricter threshold of 85% sequence similarity to an influenza cluster, Model 2 precision increased to 100% for one V gene, IGHV1-69, which has been prominently reported in antibodies reactive to influenza virus (*128*, *131*, *133*, *146–148*) (**fig. S17, C** and **D**). Therefore, both models assign distinct scores to true influenza binding sequences from some V genes related to the disease. These results reinforce the association of binding sequences with what drives Model 2 and Model 3. Applying Model 3 specifically to the 68% of known binder sequences that Model 2 was unable to score because there were no clusters with compatible V gene, J gene, and CDR3 length parameters, Model 3 achieved AUROCs averaging 0.54 +/− 0.09 (ranging up to 0.71; **fig. S17H**) and AUPRCs averaging 1.95-fold +/− 1.0 (ranging up to 3.99-fold) change over baseline prevalence. Combining the two modeling approaches, along with incorporating training data labeled at the sequence level, may therefore help guide antigen-specific antibody sequence discovery efforts.

We also evaluated sequence scores for TRB sequences downloaded from public databases of SARS-CoV-2 specific receptors (*79*). This known binder dataset was also quite small: binders made up just 0.75% of the sequence pool when combined with healthy sequences. Here, Model 2 had low performance (precision up to 1.5%, recall up to 3.9%), and there were no Model 2 clusters compatible with 99.7% of known binders, according to their TRBV gene, TBRJ gene, and CDR3 length clonal lineage parameters. This inability to evaluate most TRB binder sequences suggests Model 2’s collection of Covid-19 TCR clusters is limited, and it may explain the model’s relatively poor patient-level performance for predicting Covid-19 from TCR repertoires (**fig. S4**). According to Model 3 scores, on the other hand, TRB binder sequences were weakly favored over healthy donor sequences, with a maximum AUROC of 0.57 and a maximum AUPRC fold change of 1.36-fold in any TRBV gene.

---

**Fig. S1.**
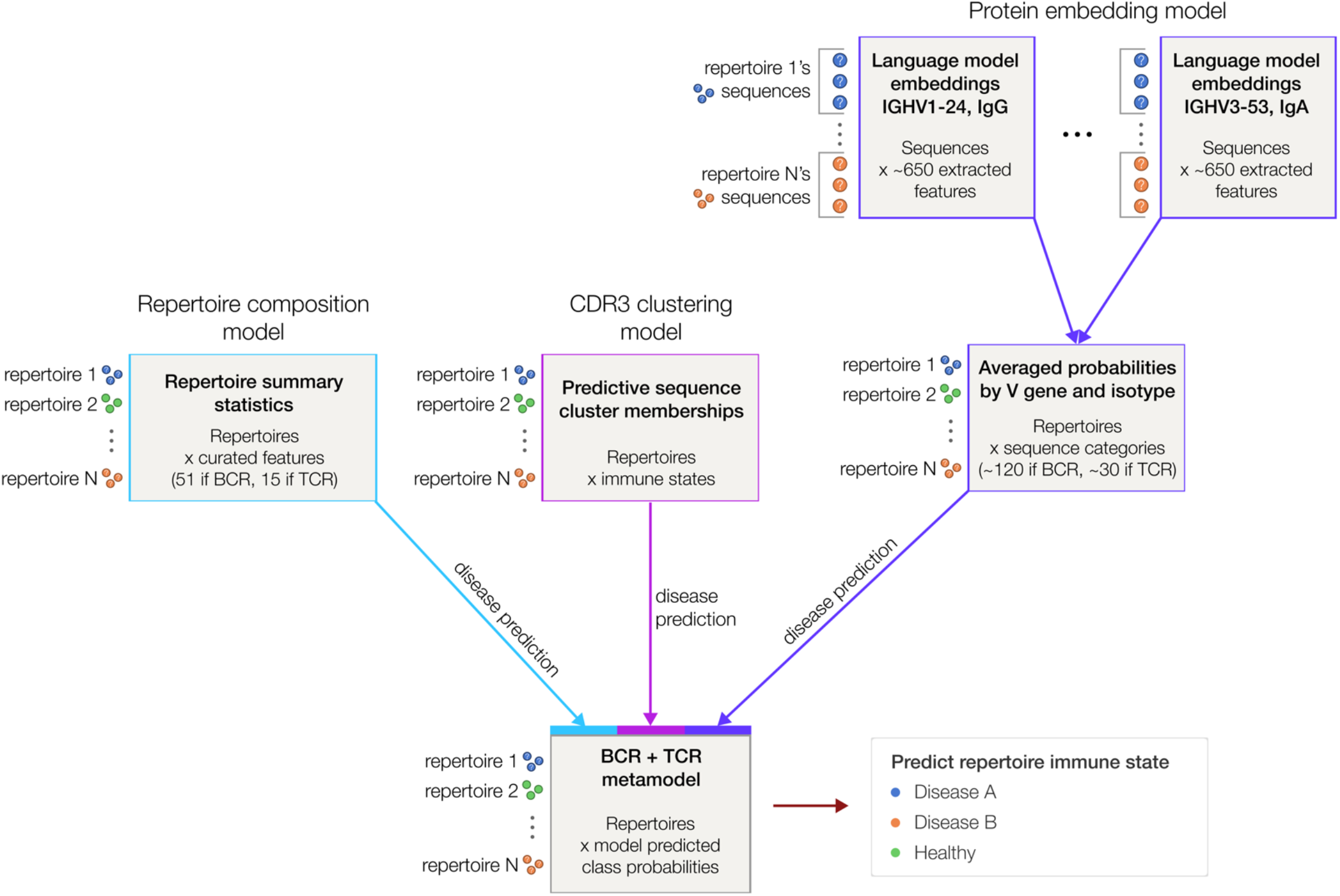
The *Mal-ID* classification pipeline for disease prediction (or other prediction tasks) has two stages. First, we fit three models per locus (i.e. three IgH and three TRB models) on a cross-validation fold’s training set. Second, we fit a metamodel on the validation set to ensemble the three inner models per locus.

**Fig. S2.**
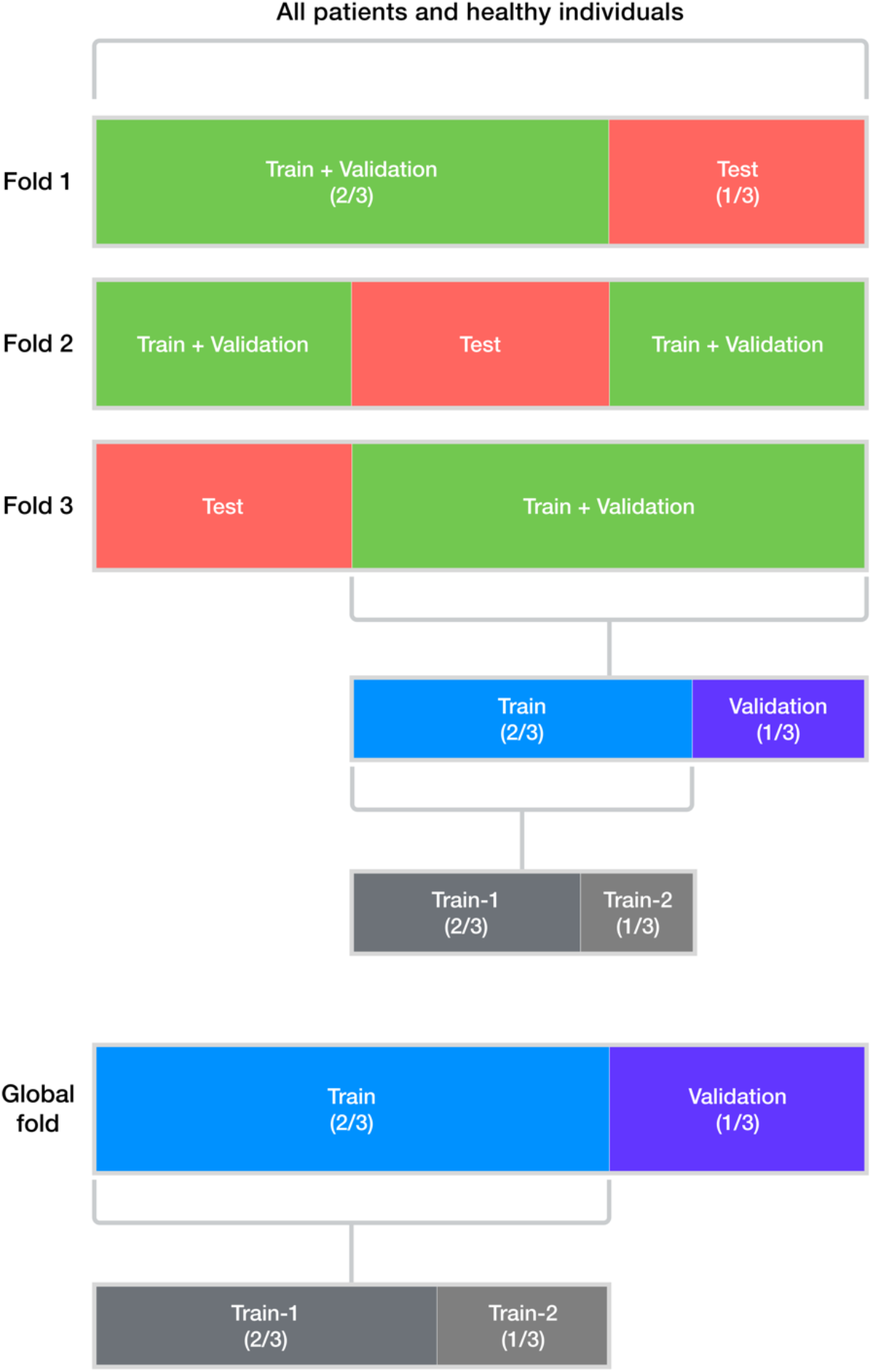
Schematic of cross-validation strategy. In each of three folds, individuals are divided into a train, validation, and test set; the training set is further divided into two pieces. All sequences from an individual are only in the train, only in the validation, or only in the test set. We also created a “global fold” to train a final model on the entire dataset, for downstream evaluation on independent cohorts.

**Fig. S3.**
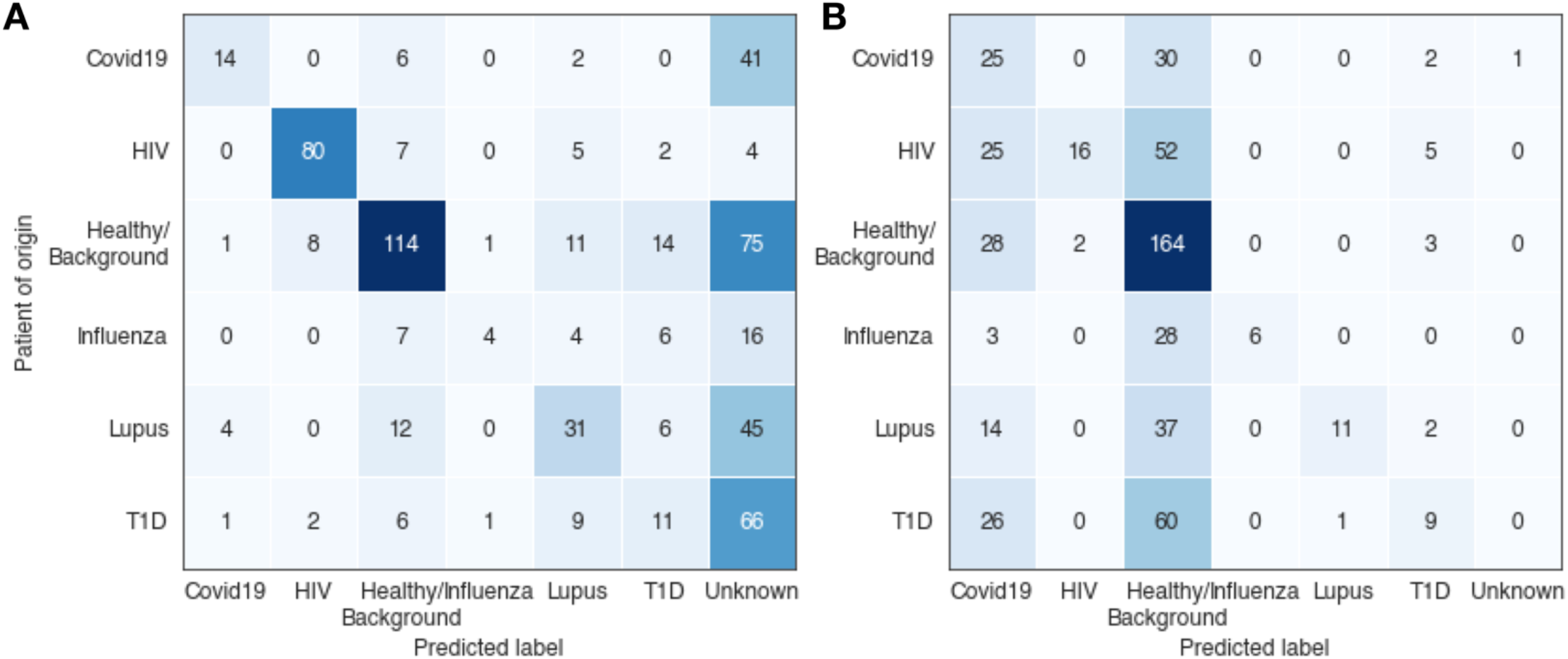
Exact CDR3 match (Emerson *et al.*, 2017) performance on held-out test set data in three cross-validation folds. (**A**) IgH repertoires; (**B**) TRB repertoires.

**Fig. S4.**
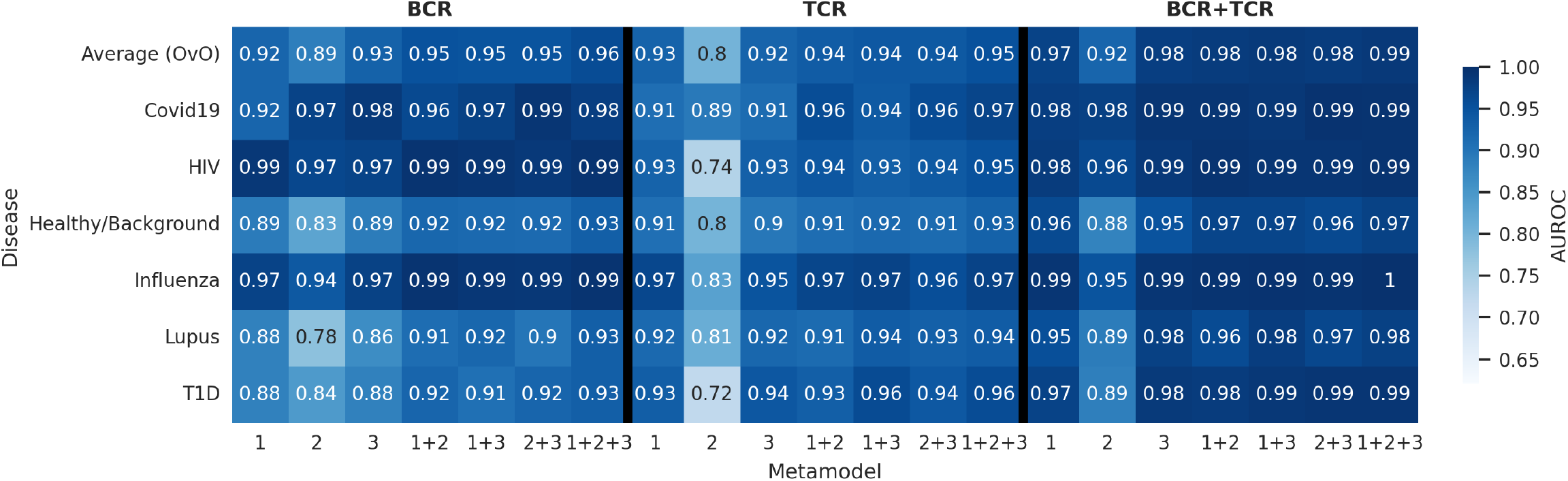
Disease classification performance by individual base models, or by ensembles of combinations of base models. Performance is summarized as AUROC scores for each class versus the rest and described by featurization methods and whether the features were extracted from BCR information, TCR information, or both. Model 1 refers to the repertoire composition classifier; model 2 refers to the CDR3 clustering classifier; and model 3 refers to the language model classifier. The CDR3 clustering models abstain from prediction on some samples, while the other models do not abstain; to create comparable metrics, these abstentions were forcibly applied to the other models. The BCR-only results include performance on BCR-only patient cohorts not present in TCR-only or BCR+TCR evaluation. The “Average (OvO)” row includes average multi-class One-vs-One (OvO) AUROC scores across classes, corresponding to the primary metric reported in this study.

**Fig. S5.**
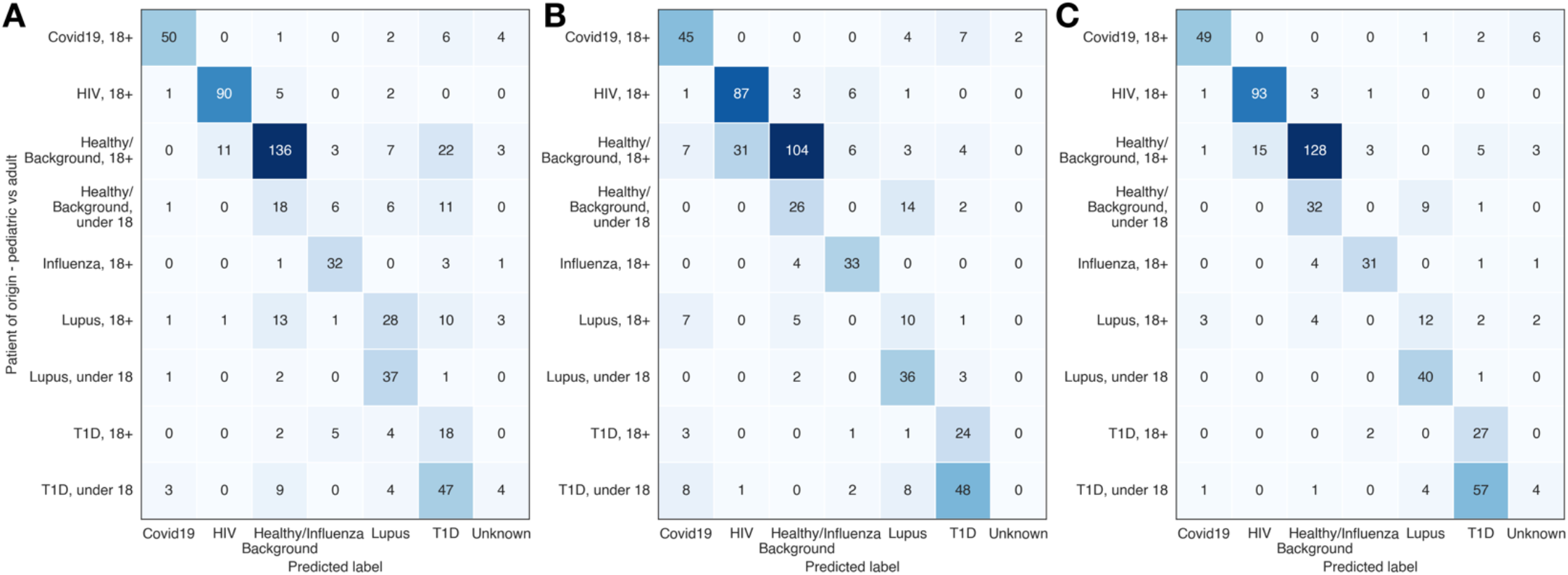
*Mal-ID* successfully differentiates between pediatric samples of different immune states. Metamodel classification performance, delineated by the ground truth disease status and age of each sample, for: (**A**) BCR-only metamodel; (**B**) TCR-only metamodel; (**C**) BCR + TCR metamodel. (The BCR and TCR metamodels have a different total number of samples due to BCR-only cohorts.)

**Fig. S6.**
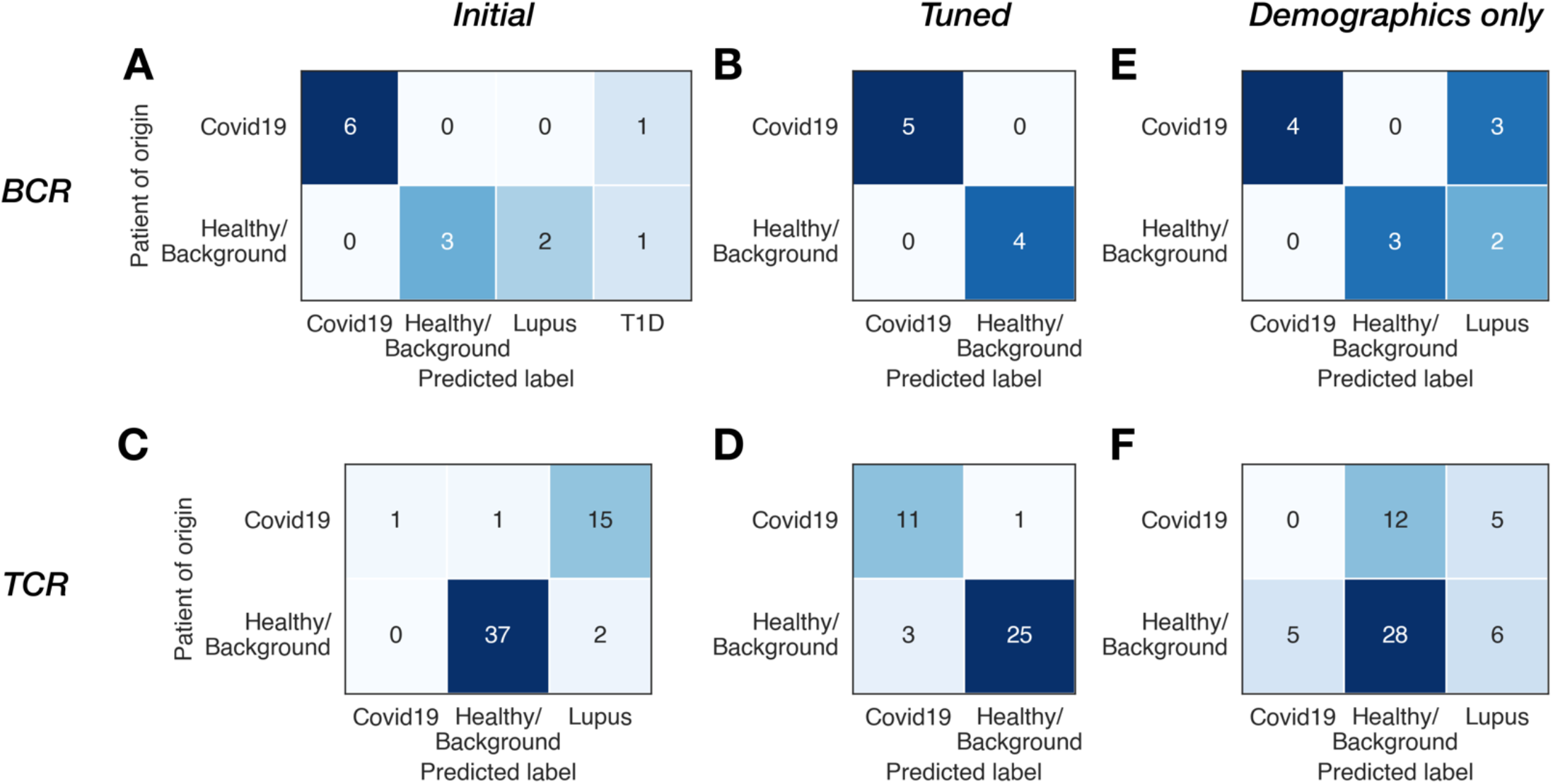
External validation cohort performance. *Mal-ID* ridge logistic regression metamodel performance on BCR or TCR external validation cohorts out of the box (first column), or with decision threshold tuning for new base rates of disease (second column). For comparison, the performance of a classifier based only on demographic information is plotted in the third column (ridge logistic regression for BCR and lasso logistic regression for TCR, which had highest performance for predicting disease from demographics in our primary dataset). One healthy BCR sample was not evaluated by the demographics-only classifier because it did not have complete demographic information.

**Fig. S7.**
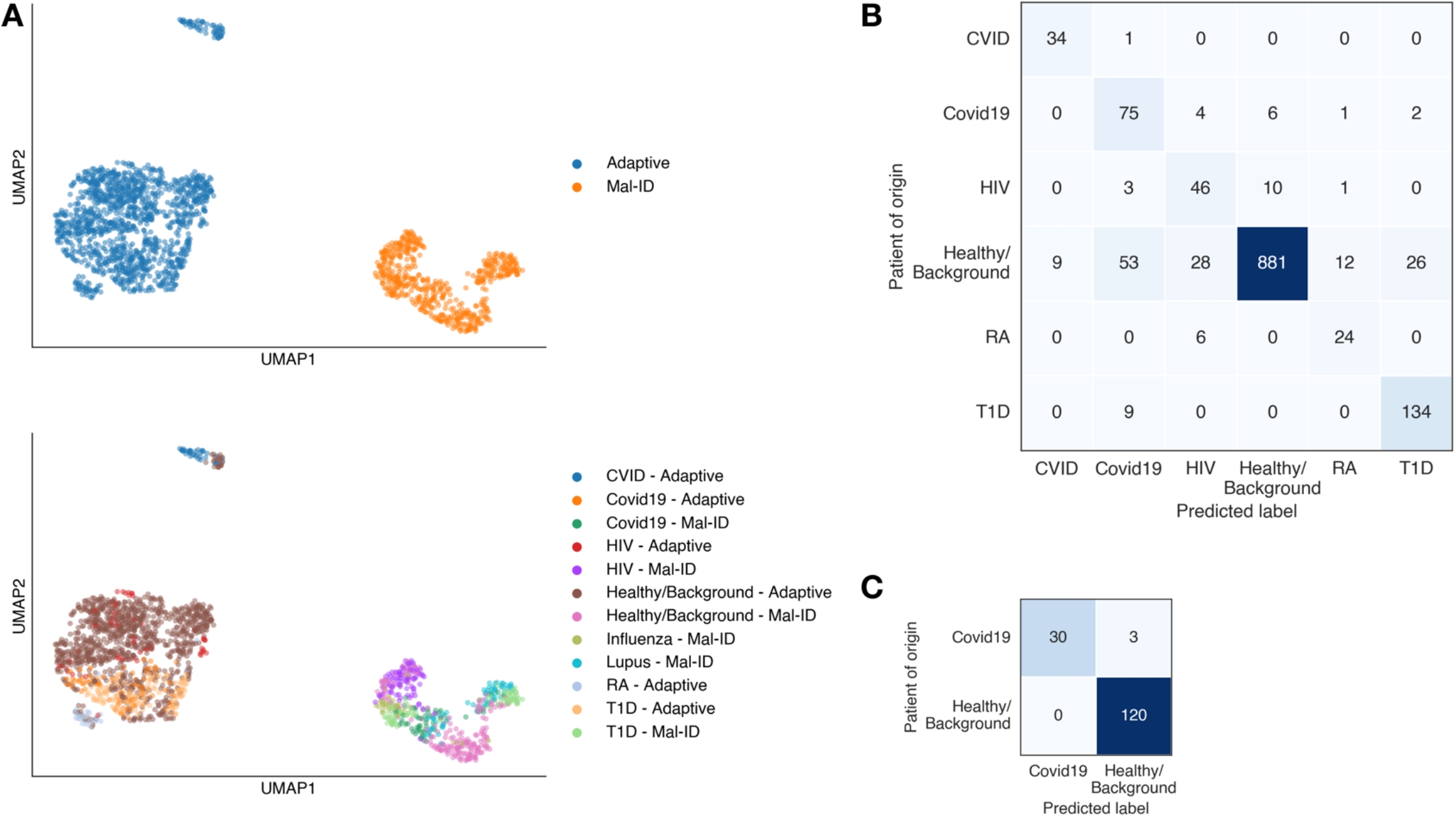
*Mal-ID* framework can be applied to genomic DNA sequencing data. (**A**) External cohorts from Adaptive Biotechnologies sequencing have different V gene usage than the *Mal-ID* cDNA sequencing primary dataset. A UMAP of TRBV gene use proportions by sample (excluding rare V genes, to avoid disproportionate effects from minute differences in their proportions) shows that Adaptive cohort V gene use is systematically different from our cohorts. (**B**) Classification results from training the *Mal-ID* architecture on TRB-only Adaptive Biotechnologies data. Three cross-validation folds shown, including six classes: common variable immunodeficiency (CVID), Covid-19, HIV, rheumatoid arthritis (RA), type-1 diabetes (T1D), and healthy. (**C**) *Mal-ID* was also retrained on only the data from a healthy study and a Covid-19 study that both included several cohorts (*17*, *79*). Entire Covid-19 and healthy cohorts were held-out from training and used only for evaluation. Dataset descriptions are provided in **table S4**.

**Fig. S8.**
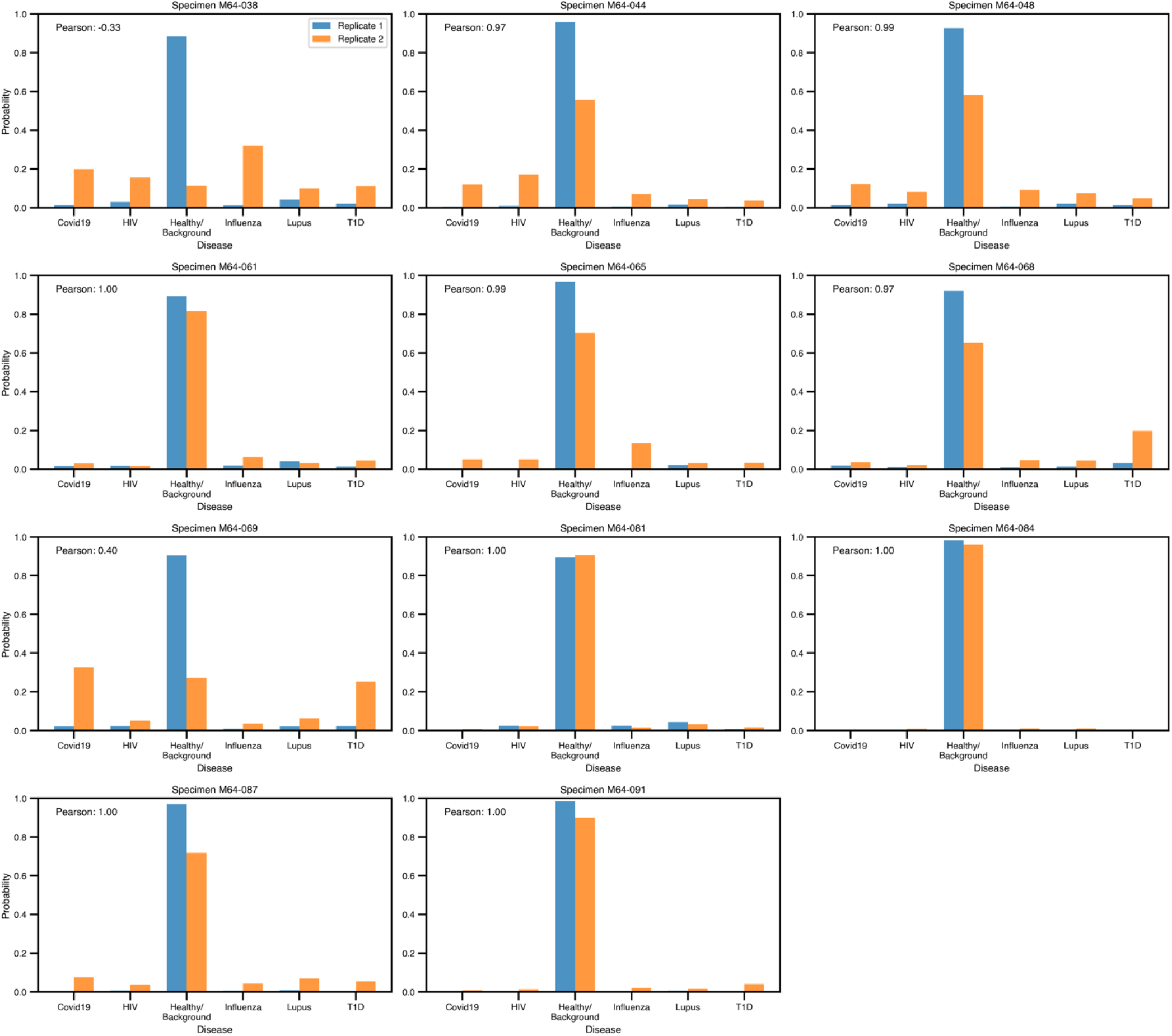
Consistent predictions for replicate datasets. Predicted class probabilities for original sample (blue) and re-sequenced replicate sample (orange) of 11 healthy donors, all held out from the training process. Correlation scores indicate high similarity between the predicted class probability vectors for the two replicates of most samples.

**Fig. S9.**
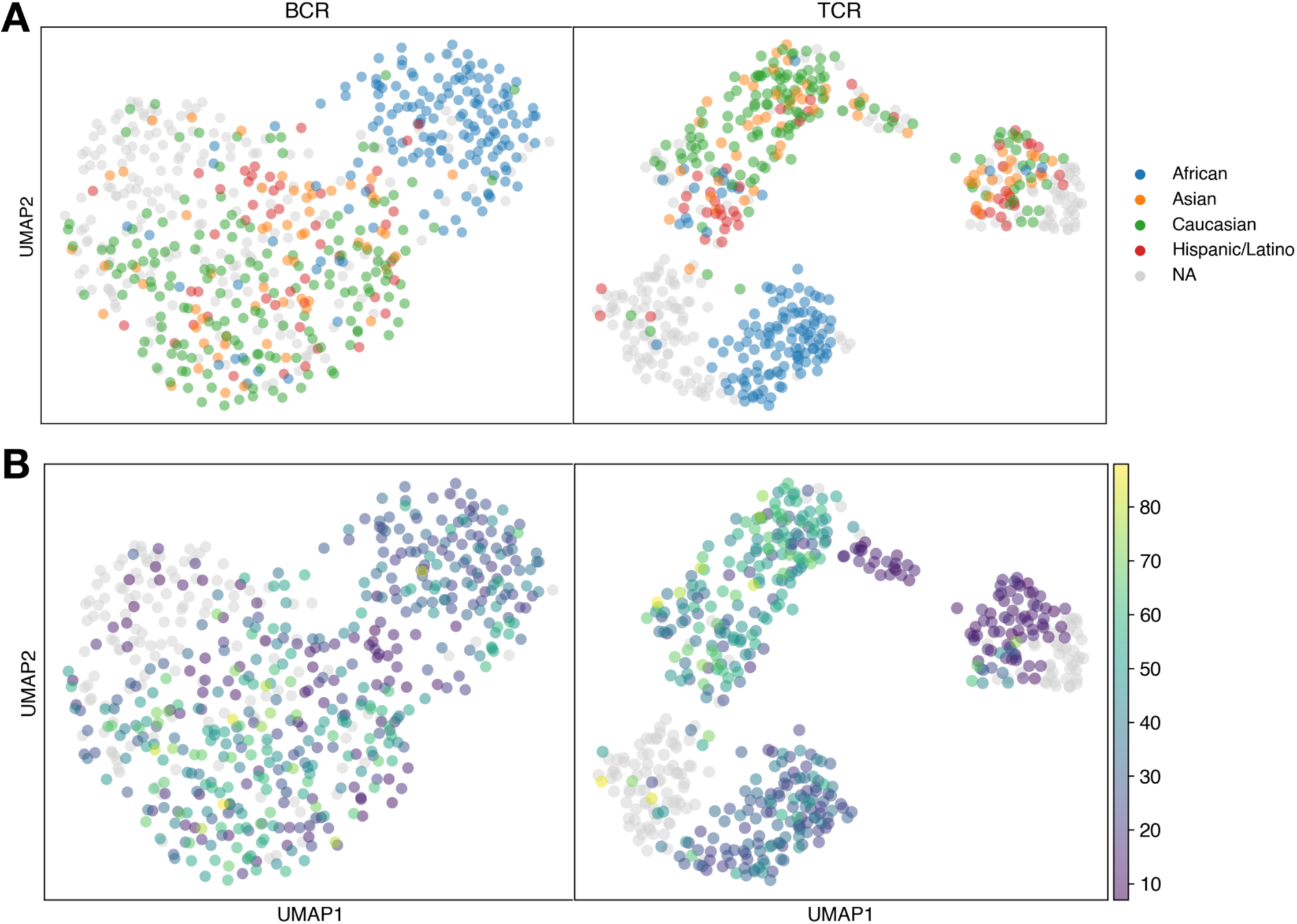
Demographic traits appear related to V gene usage trends. V gene usage proportions of IgH (left panels) and TRB (right panels) repertoires in the *Mal-ID* dataset, visualized with UMAP and colored by (**A**) ancestry or (**B**) age. (Rare V genes are excluded, as in *Mal-ID* models 1 and 3.)

**Fig. S10.**
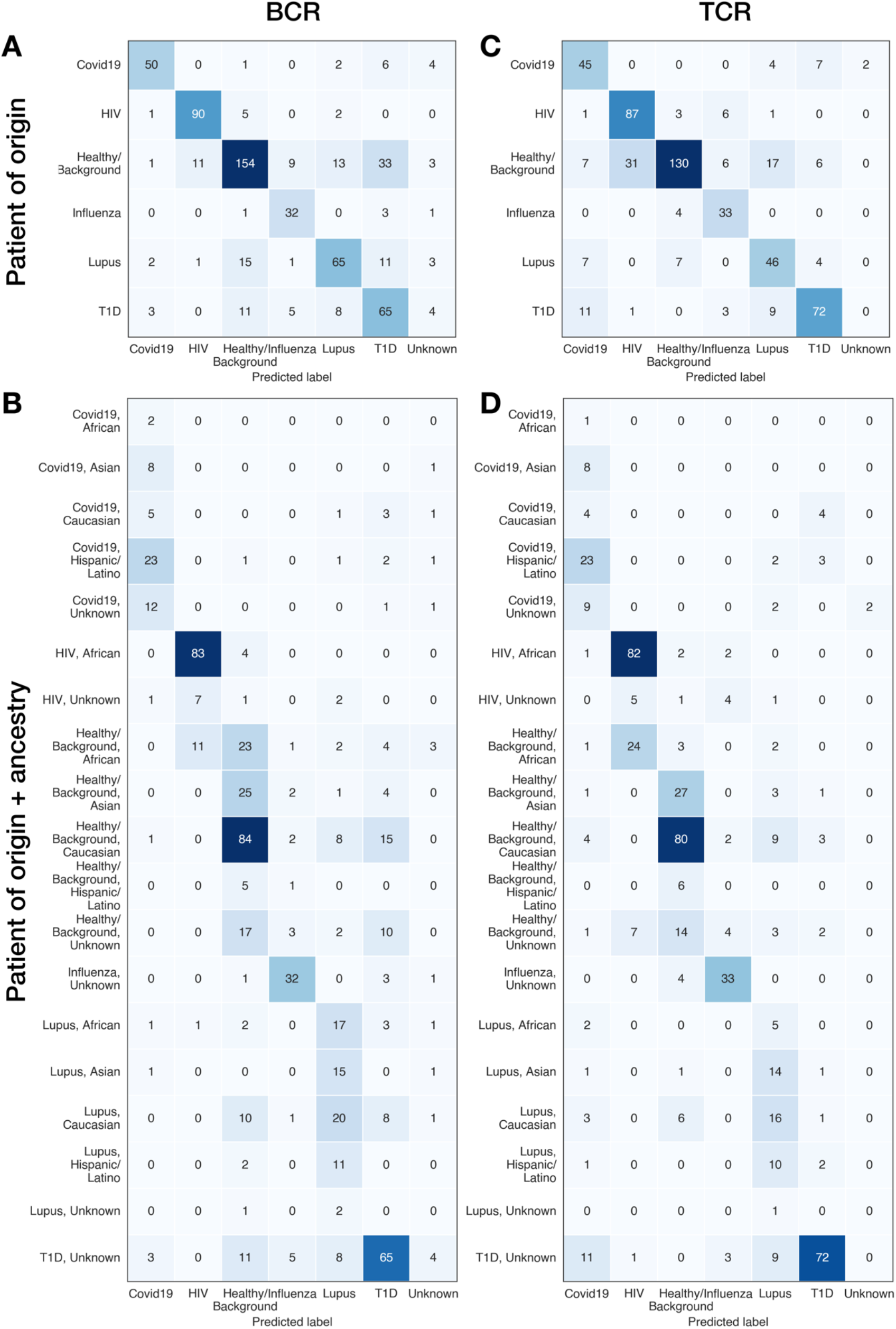
BCR-only and TCR-only ensemble models show differences in disease classification. Delineating by the ground truth disease status and ancestry of each sample (B and D) shows that the “Healthy/Background, African” cohort, a healthy control group corresponding to the HIV cohort and whose members are predominantly African and live in Africa, is misclassified as HIV by the TCR model (**C** and **D**), but not by the BCR model (**A** and **B**). (The BCR and TCR metamodels have a different total number of samples due to BCR-only cohorts.)

**Fig. S11.**
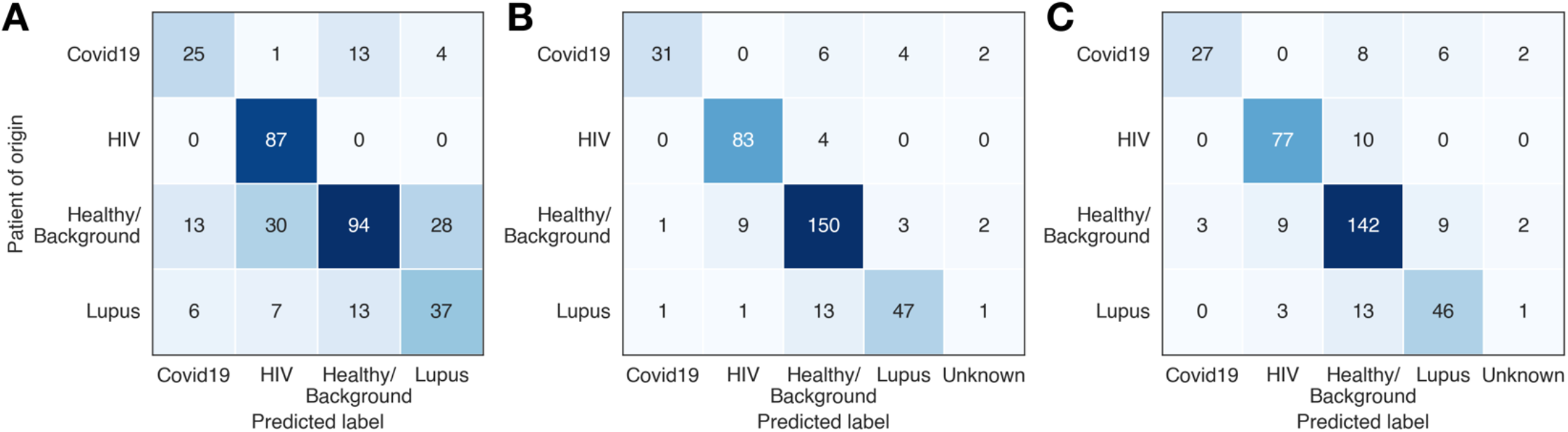
Demographic covariates have limited impact on disease classification. (**A**) Metamodel classification performance using only age, sex, and ancestry features, without any sequence features. (**B**) Metamodel classification performance using age, sex, and ancestry demographic features, along with sequence features, and interaction terms between these two sets of features. (**C**) Metamodel classification performance using sequence features only, with age, sex, and ancestry regressed out.

**Fig. S12.**
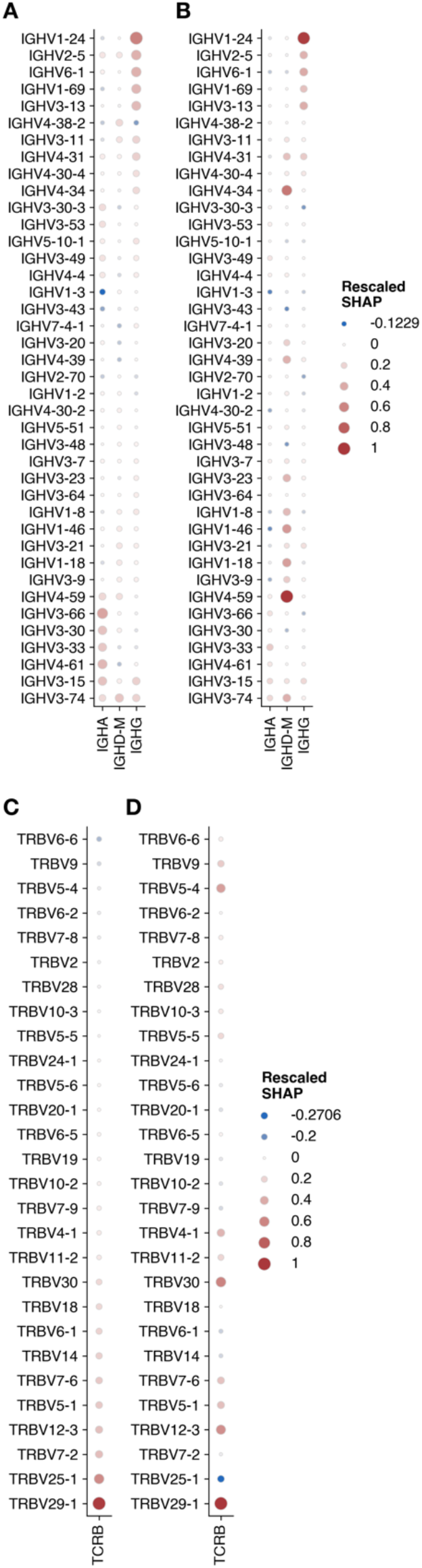
Signal contributing to lupus prediction can be derived from different sequence categories, depending on the patient. SHAP values for the lupus-vs-rest Model 3 aggregation step were clustered, with average (rescaled) SHAP values plotted for each cluster of positive examples. BCR: (**A**) cluster 1 is 73% adult; (**B**) cluster 2 is 64% adult. TCR: (**C**) cluster 1 is 88% adult; (**D**) cluster 2 is 100% pediatric.

**Fig. S13.**
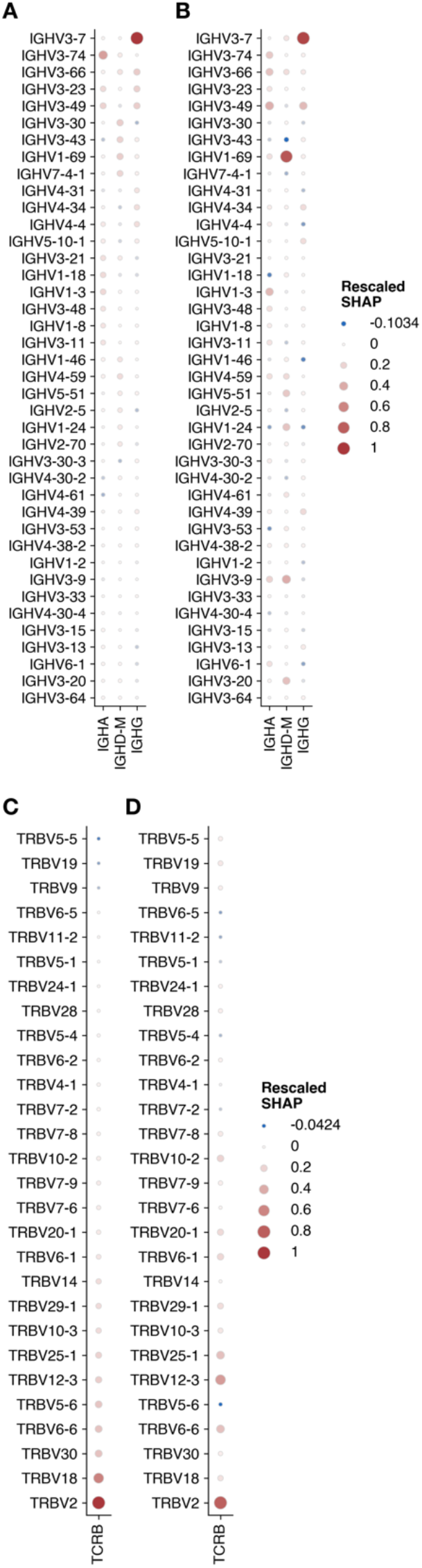
Signal contributing to T1D prediction can be derived from different sequence categories, depending on the patient. SHAP values for the T1D-vs-rest Model 3 aggregation step were clustered, with average SHAP values plotted for each cluster of positive examples. BCR: (**A**) cluster 1 is 64% pediatric; (**B**) cluster 2 is 63% pediatric. TCR: (**C**) cluster 1 is 71% pediatric; (**D**) cluster 2 is 50% pediatric.

**Fig. S14.**
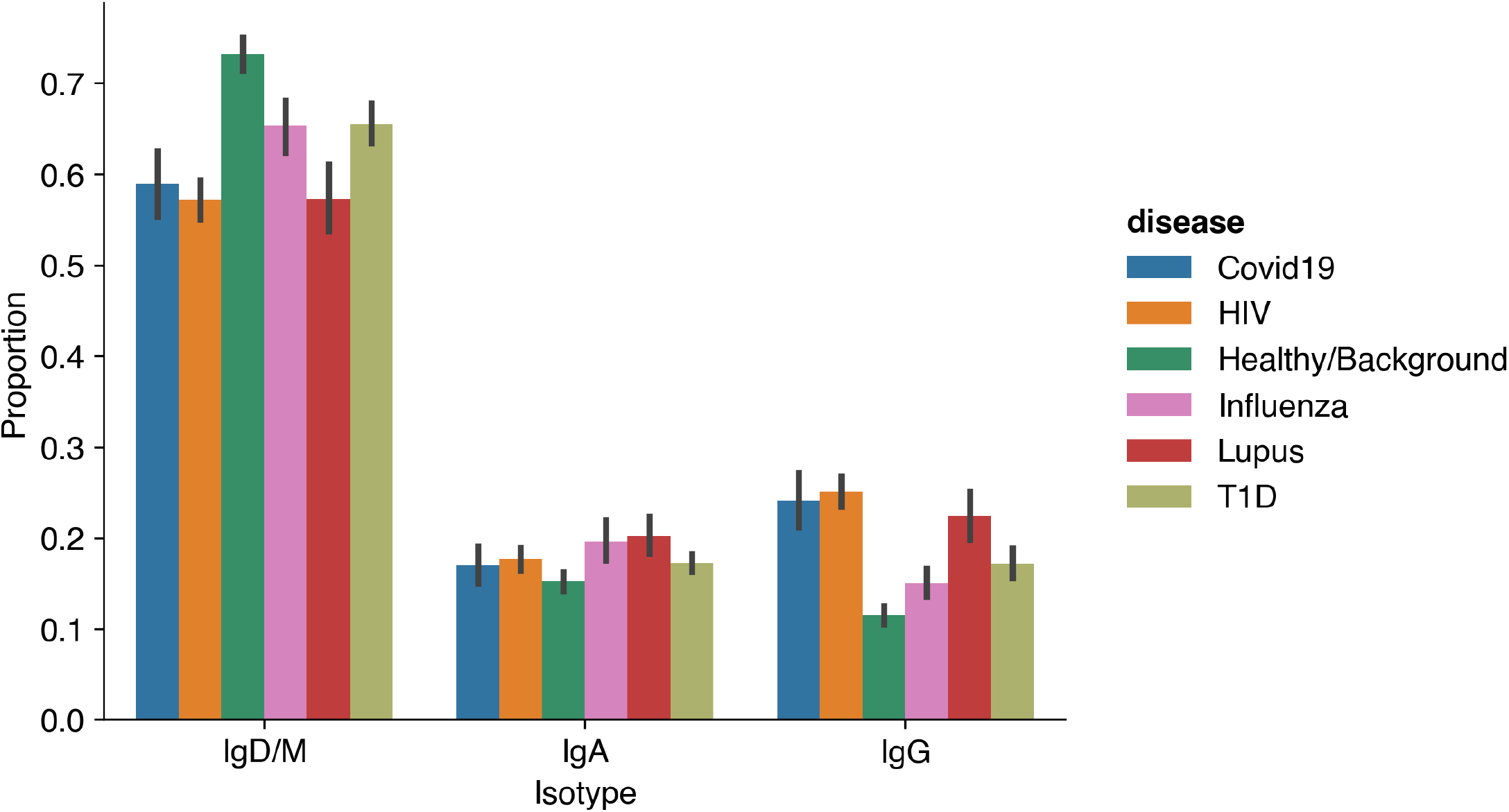
Isotype proportions per sample present in the IgH data are different between immune states. Average and 95% confidence interval are shown. Differences in isotype proportions are technical artifacts, so the models are designed to not quantify isotype proportions. Instead, Models 1 and 3 evaluate each isotype separately, and Model 2 uses all the data jointly without regard for isotype.

**Fig. S15.**
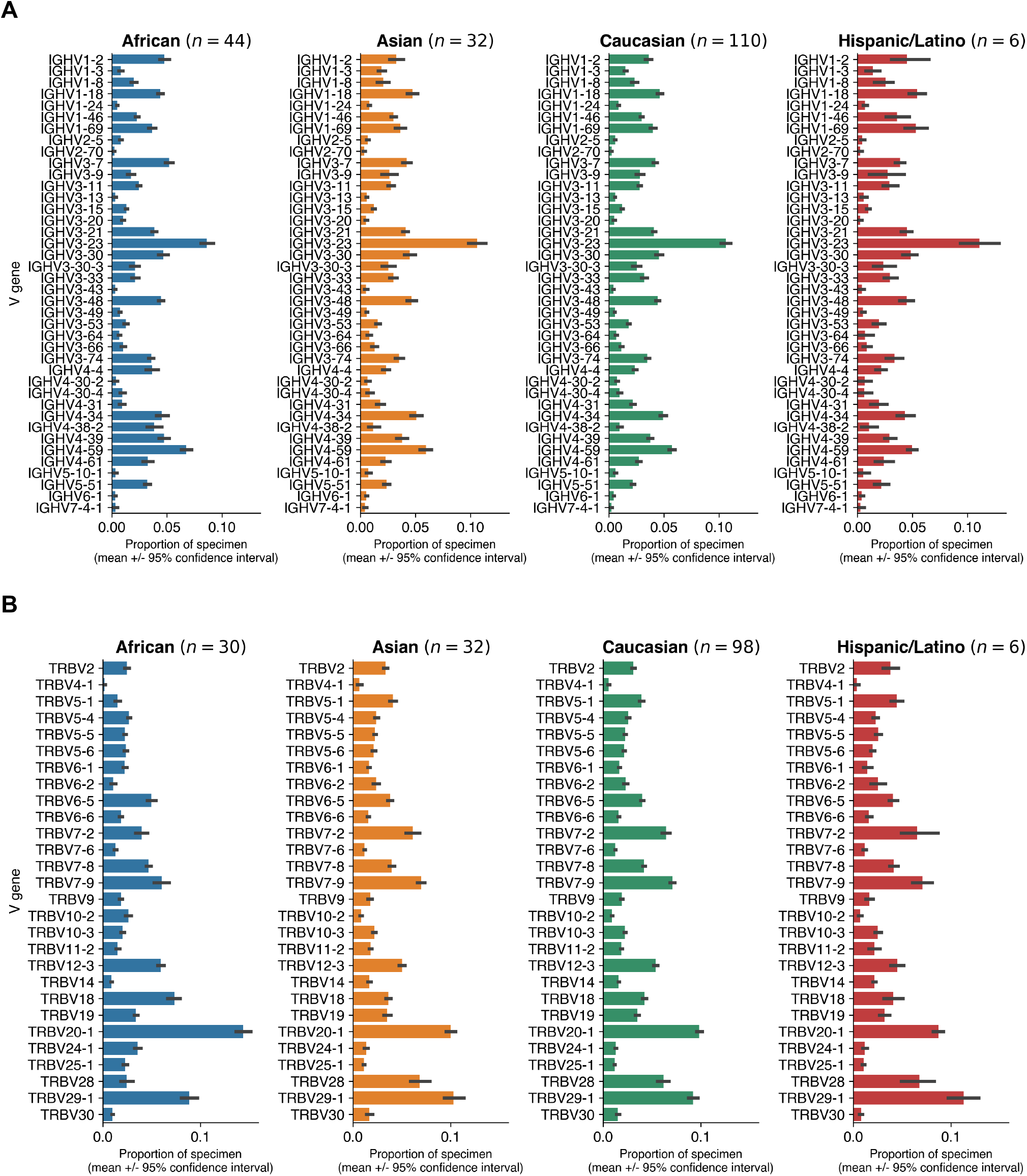
Some V gene usage frequencies appear related to ancestry. IGHV or TRBV gene use proportions in healthy control samples, stratified by ancestry, with average and 95% confidence interval plotted. (**A**) BCR (note higher sample sizes due to presence of BCR-only cohorts). (**B**) TCR.

**Fig. S16.**
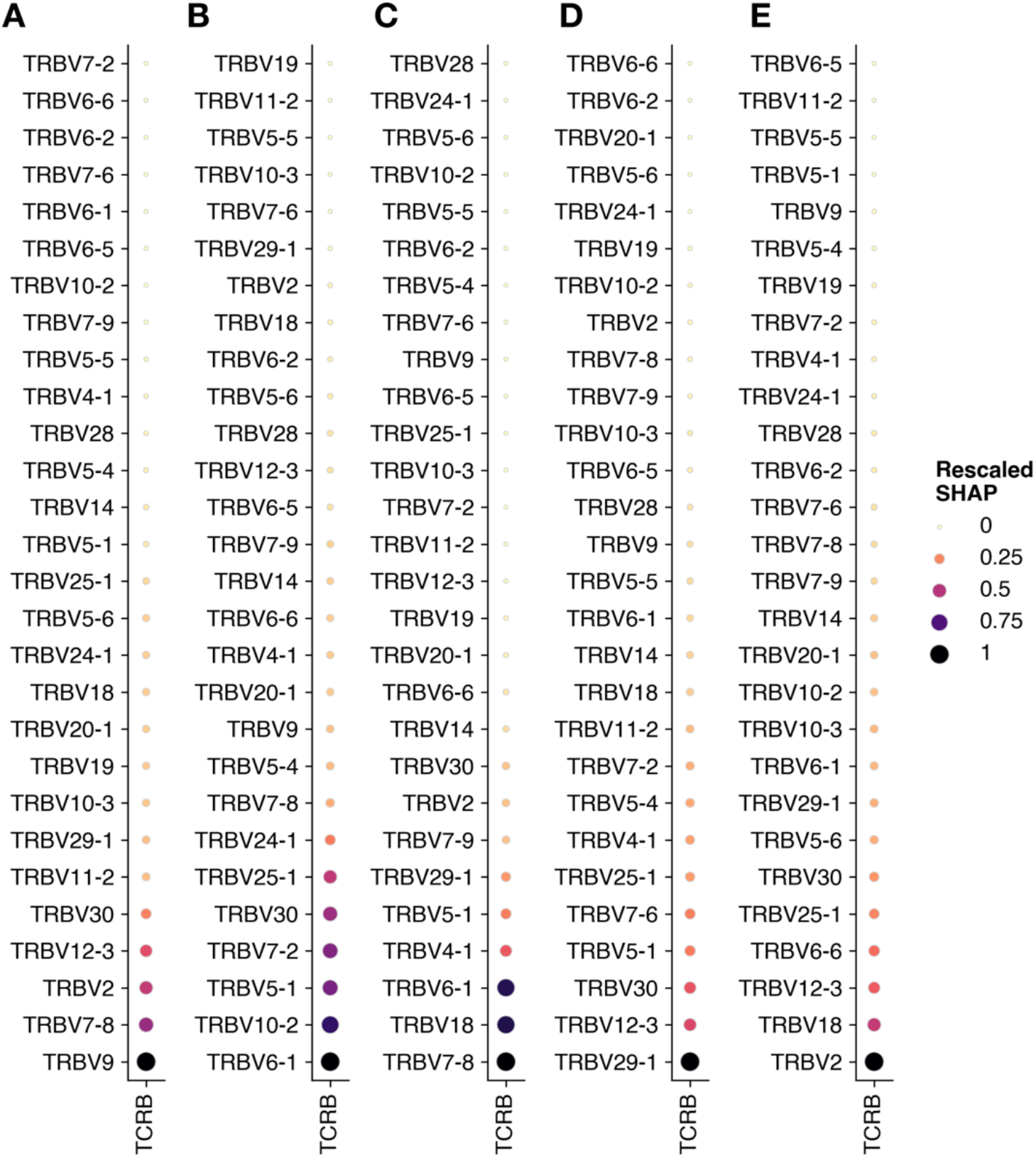
Disease-associated TRBV genes prioritized by Model 3 using protein language embeddings. Shapley importance (SHAP) values quantifying the contribution of average sequence predictions from each TRBV gene category to Model 3’s prediction of a sample’s disease state are plotted for (**A**) Covid-19, (**B**) HIV, (**C**) influenza vaccination, (**D**) lupus; and (**E**) type-1 diabetes.

**Fig. S17.**
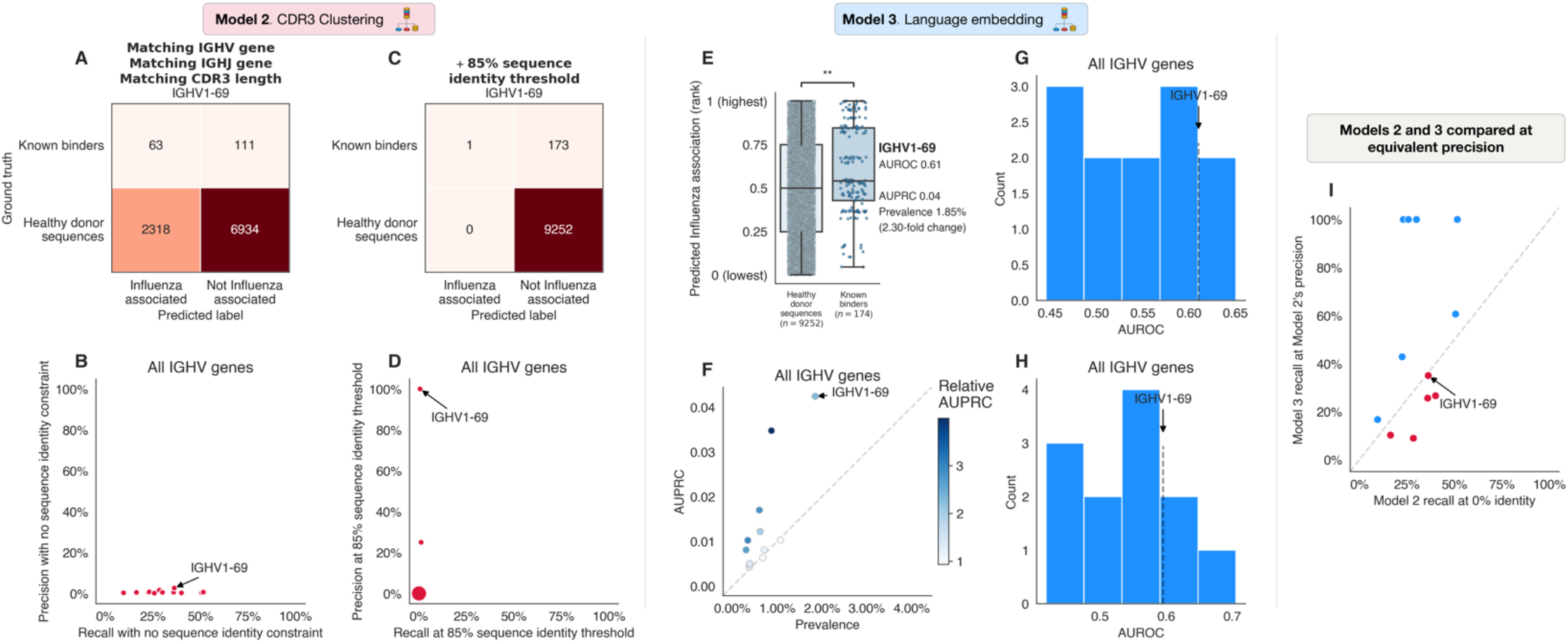
Models 2 and 3 learn influenza antigen-specific sequence patterns from influenza vaccine recipient data and can distinguish between known influenza-specific antibody sequences and healthy donor sequences. For this comparison, validated SARS-CoV-2-binding sequences (*78*) and a subset of healthy donor sequences were held out from training. Known binder detection using Model 2 or Model 3 predictions of sequence association to disease was evaluated separately for each IGHV gene; performance is shown for IGHV1-69 and compared across IGHV genes. **(A** to **D)** Model 2 identifies a conservative set of public clones enriched in influenza vaccine recipients which match some known binders. Model 2’s precision and recall across IGHV genes is shown, with binding predictions determined: (**A** and **B**) based on shared IGHV gene, IGHJ gene, and CDR3 length with any influenza cluster identified in Model 2’s training procedure; or (**C** and **D**) at an 85% CDR3 sequence identity threshold. (**E** to **H**) Model 3 ranks known binders higher than healthy sequences based on predicted influenza probability (**E**), with relative AUPRC ranging up to 4.0-fold over baseline prevalence (**F**) and AUROC up to 0.65 across IGHV genes (**G**). Permutation test in panel (E) to assess whether IGHV1-24 known binders have higher ranks than healthy donor sequences, with consistent labels maintained during the permutation process across sequences from each healthy donor: p value 0.008. (**H**) Model 3 maintains reasonable performance (AUROC up to 0.71) for sequences that are not evaluated by Model 2’s clustering (sequences for which Model 2 identified no SARS-CoV-2 clusters with matching IGHV gene, IGHJ gene, and CDR3 length). (**I**) At equivalent precision, recall was not consistently higher for Model 2 or Model 3. IGHV genes where Model 3 has higher recall than Model 2 are shown in blue. For each IGHV gene, recall was calculated for Models 2 and 3 at Model 2’s precision shown in (B). Point size indicates number of identical values plotted at a particular location for panels (B), (D), and (I).

**Table S1:**
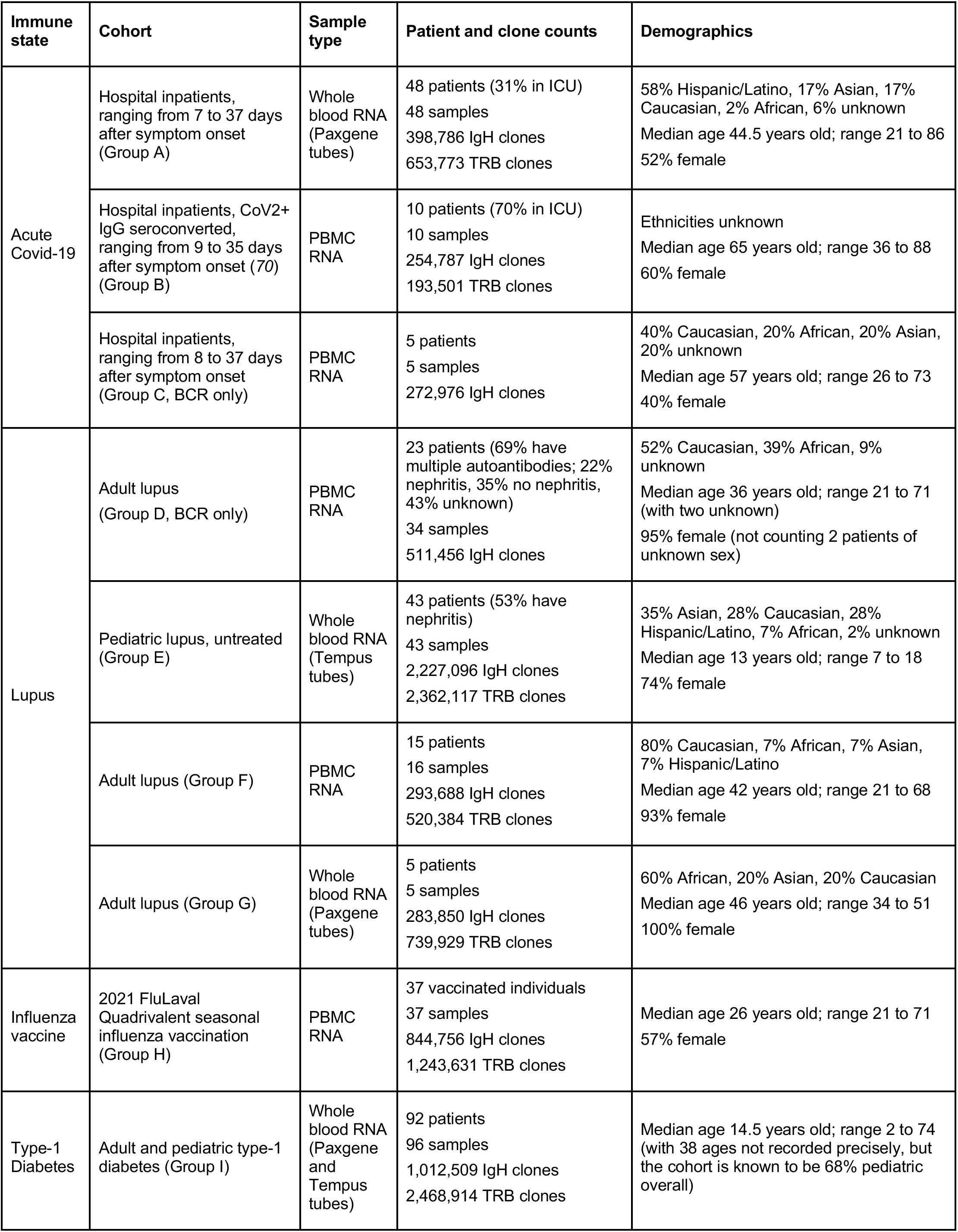
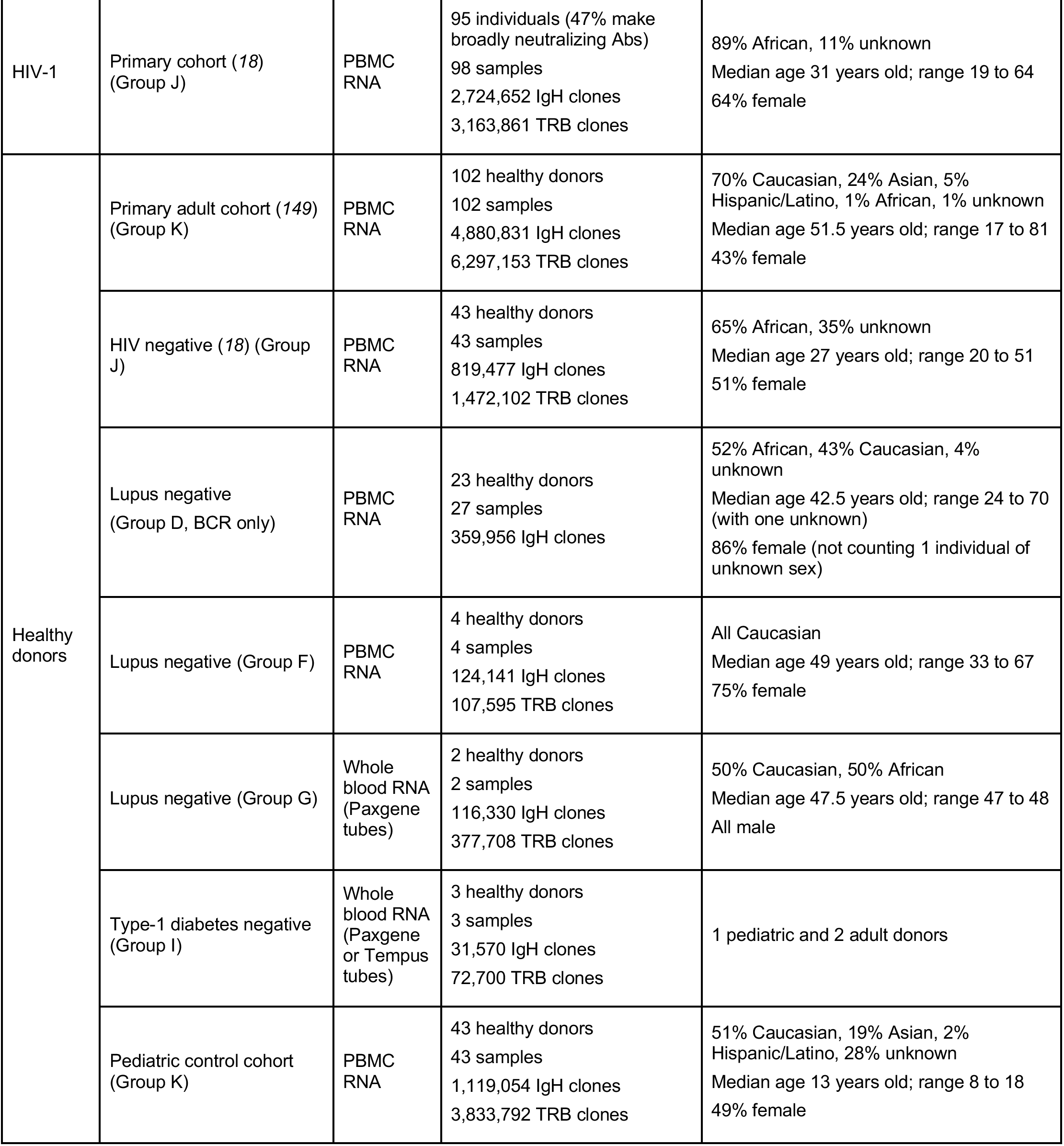
Cohort and batch info for 593 individuals with a total of 616 samples. 550 of the 616 samples had both BCR and TCR sequencing performed, representing 542 of the total 593 individuals. The remainder only underwent BCR IgH sequencing. The alphabetical Group labels indicate which samples were processed together.

| Immune state | Cohort | Sample type | Patient and clone counts | Demographics |
| --- | --- | --- | --- | --- |
| Acute Covid-19 | Hospital inpatients, ranging from 7 to 37 days after symptom onset (Group A) | Whole blood RNA (Paxgene tubes) | 48 patients (31% in ICU)<br>48 samples<br>398,786 IgH clones<br>653,773 TRB clones | 58% Hispanic/Latino, 17% Asian, 17% Caucasian, 2% African, 6% unknown<br>Median age 44.5 years old; range 21 to 86<br>52% female |
|  | Hospital inpatients, CoV2+ IgG seroconverted, ranging from 9 to 35 days after symptom onset (70) (Group B) | PBMC RNA | 10 patients (70% in ICU)<br>10 samples<br>254,787 IgH clones<br>193,501 TRB clones | Ethnicities unknown<br>Median age 65 years old; range 36 to 88<br>60% female |
|  | Hospital inpatients, ranging from 8 to 37 days after symptom onset (Group C, BCR only) | PBMC RNA | 5 patients<br>5 samples<br>272,976 IgH clones | 40% Caucasian, 20% African, 20% Asian, 20% unknown<br>Median age 57 years old; range 26 to 73<br>40% female |
| Lupus | Adult lupus (Group D, BCR only) | PBMC RNA | 23 patients (69% have multiple autoantibodies; 22% nephritis, 35% no nephritis, 43% unknown)<br>34 samples<br>511,456 IgH clones | 52% Caucasian, 39% African, 9% unknown<br>Median age 36 years old; range 21 to 71 (with two unknown)<br>95% female (not counting 2 patients of unknown sex) |
|  | Pediatric lupus, untreated (Group E) | Whole blood RNA (Tempus tubes) | 43 patients (53% have nephritis)<br>43 samples<br>2,227,096 IgH clones<br>2,362,117 TRB clones | 35% Asian, 28% Caucasian, 28% Hispanic/Latino, 7% African, 2% unknown<br>Median age 13 years old; range 7 to 18<br>74% female |
|  | Adult lupus (Group F) | PBMC RNA | 15 patients<br>16 samples<br>293,688 IgH clones<br>520,384 TRB clones | 80% Caucasian, 7% African, 7% Asian, 7% Hispanic/Latino<br>Median age 42 years old; range 21 to 68<br>93% female |
|  | Adult lupus (Group G) | Whole blood RNA (Paxgene tubes) | 5 patients<br>5 samples<br>283,850 IgH clones<br>739,929 TRB clones | 60% African, 20% Asian, 20% Caucasian<br>Median age 46 years old; range 34 to 51<br>100% female |
| Influenza vaccine | 2021 FluLaval Quadrivalent seasonal influenza vaccination (Group H) | PBMC RNA | 37 vaccinated individuals<br>37 samples<br>844,756 IgH clones<br>1,243,631 TRB clones | Median age 26 years old; range 21 to 71<br>57% female |
| Type-1 Diabetes | Adult and pediatric type-1 diabetes (Group I) | Whole blood RNA (Paxgene and Tempus tubes) | 92 patients<br>96 samples<br>1,012,509 IgH clones<br>2,468,914 TRB clones | Median age 14.5 years old; range 2 to 74 (with 38 ages not recorded precisely, but the cohort is known to be 68% pediatric overall) |
| HIV-1 | Primary cohort (18)<br>(Group J) | PBMC<br>RNA | 95 individuals (47% make broadly neutralizing Abs)<br>98 samples<br>2,724,652 IgH clones<br>3,163,861 TRB clones | 89% African, 11% unknown<br>Median age 31 years old; range 19 to 64<br>64% female |
| Healthy donors | Primary adult cohort (149)<br>(Group K) | PBMC<br>RNA | 102 healthy donors<br>102 samples<br>4,880,831 IgH clones<br>6,297,153 TRB clones | 70% Caucasian, 24% Asian, 5% Hispanic/Latino, 1% African, 1% unknown<br>Median age 51.5 years old; range 17 to 81<br>43% female |
|  | HIV negative (18) (Group J) | PBMC<br>RNA | 43 healthy donors<br>43 samples<br>819,477 IgH clones<br>1,472,102 TRB clones | 65% African, 35% unknown<br>Median age 27 years old; range 20 to 51<br>51% female |
|  | Lupus negative<br>(Group D, BCR only) | PBMC<br>RNA | 23 healthy donors<br>27 samples<br>359,956 IgH clones | 52% African, 43% Caucasian, 4% unknown<br>Median age 42.5 years old; range 24 to 70 (with one unknown)<br>86% female (not counting 1 individual of unknown sex) |
|  | Lupus negative (Group F) | PBMC<br>RNA | 4 healthy donors<br>4 samples<br>124,141 IgH clones<br>107,595 TRB clones | All Caucasian<br>Median age 49 years old; range 33 to 67<br>75% female |
|  | Lupus negative (Group G) | Whole blood RNA<br>(Paxgene tubes) | 2 healthy donors<br>2 samples<br>116,330 IgH clones<br>377,708 TRB clones | 50% Caucasian, 50% African<br>Median age 47.5 years old; range 47 to 48<br>All male |
|  | Type-1 diabetes negative<br>(Group I) | Whole blood RNA<br>(Paxgene or Tempus tubes) | 3 healthy donors<br>3 samples<br>31,570 IgH clones<br>72,700 TRB clones | 1 pediatric and 2 adult donors |
|  | Pediatric control cohort<br>(Group K) | PBMC<br>RNA | 43 healthy donors<br>43 samples<br>1,119,054 IgH clones<br>3,833,792 TRB clones | 51% Caucasian, 19% Asian, 2% Hispanic/Latino, 28% unknown<br>Median age 13 years old; range 8 to 18<br>49% female |

**Table S2:** Average cross-validated test set performance on 616 BCR samples, 550 TCR samples, or 550 BCR + TCR samples using ensemble metamodels trained with individual model components or combinations thereof. AUPRC stands for area under the precision-recall curve. Abstentions hurt accuracy scores (they count as incorrect predictions), but are not included in the calculation of probability-based metrics AUROC and AUPRC, because no predicted class probabilities are generated for abstained samples.

| Model Components Used | Locus | Accuracy | AUROC | AUPRC | Abstention rate |
| --- | --- | --- | --- | --- | --- |
| Model 1:<br>Global repertoire statistics | BCR | 70.1% | 0.919 +/- 0.007 | 0.923 +/- 0.004 | N/A |
|  | TCR | 69.8% | 0.925 +/- 0.008 | 0.921 +/- 0.010 | N/A |
|  | BCR + TCR | 81.8% | 0.971 +/- 0.003 | 0.971 +/- 0.003 | N/A |
| Model 2:<br>CDR3 sequence clustering | BCR | 59.6% | 0.887 +/- 0.016 | 0.889 +/- 0.010 | 2.4% |
|  | TCR | 51.1% | 0.802 +/- 0.024 | 0.806 +/- 0.022 | 0.4% |
|  | BCR + TCR | 67.3% | 0.922 +/- 0.012 | 0.924 +/- 0.009 | 2.9% |
| Model 3:<br>Language model embedding | BCR | 66.4% | 0.923 +/- 0.012 | 0.921 +/- 0.014 | N/A |
|  | TCR | 68.7% | 0.924 +/- 0.026 | 0.917 +/- 0.027 | N/A |
|  | BCR + TCR | 82.0% | 0.975 +/- 0.010 | 0.975 +/- 0.010 | N/A |
| Models 1 + 2 | BCR | 72.4% | 0.950 +/- 0.005 | 0.950 +/- 0.005 | 2.4% |
|  | TCR | 71.6% | 0.936 +/- 0.007 | 0.931 +/- 0.009 | 0.4% |
|  | BCR + TCR | 81.3% | 0.981 +/- 0.001 | 0.980 +/- 0.002 | 2.9% |
| Models 1 + 3 | BCR | 73.7% | 0.944 +/- 0.007 | 0.947 +/- 0.005 | N/A |
|  | TCR | 71.8% | 0.941 +/- 0.006 | 0.934 +/- 0.006 | N/A |
|  | BCR + TCR | 84.0% | 0.982 +/- 0.005 | 0.981 +/- 0.004 | N/A |
| Models 2 + 3 | BCR | 71.3% | 0.948 +/- 0.002 | 0.945 +/- 0.004 | 2.4% |
|  | TCR | 72.5% | 0.940 +/- 0.002 | 0.933 +/- 0.006 | 0.4% |
|  | BCR + TCR | 80.7% | 0.982 +/- 0.008 | 0.981 +/- 0.008 | 2.9% |
| Models 1 + 2 + 3 | BCR | 74.0% | 0.959 +/- 0.001 | 0.959 +/- 0.002 | 2.4% |
|  | TCR | 75.1% | 0.952 +/- 0.003 | 0.943 +/- 0.008 | 0.4% |
|  | BCR + TCR<br>(Fig. 2A) | 85.3% | 0.986 +/- 0.002 | 0.986 +/- 0.002 | 2.9% |

**Table S3:**
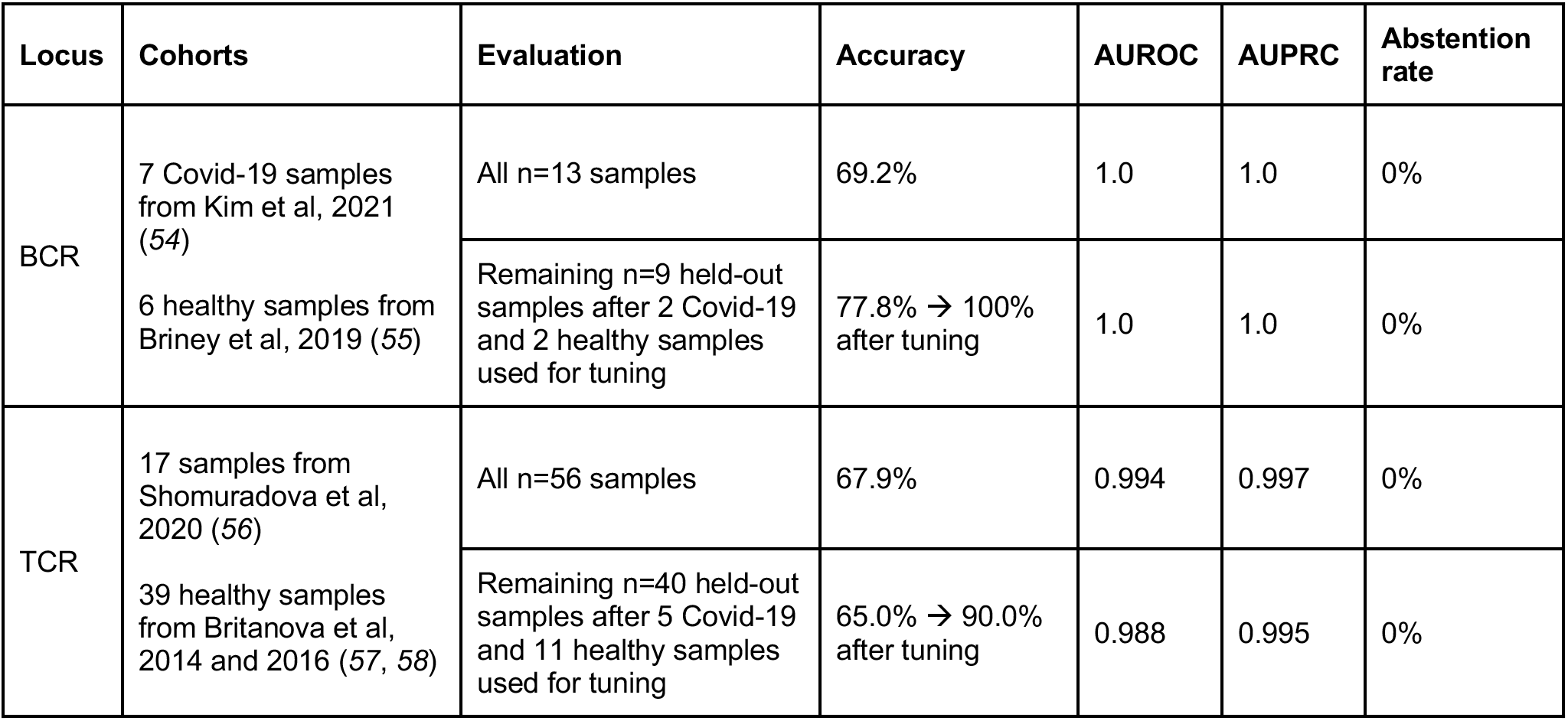
External validation cohort performance using BCR-only or TCR-only ridge logistic regression ensemble models. First, performance on the full data was reported. Then the evaluation was repeated with a portion of samples set aside for tuning class decision thresholds; the remaining samples were used for calculating performance before and after tuning.

**Table S4:** Peripheral blood genomic DNA datasets from Adaptive Biotechnologies, comprising 1365 samples, used to retrain and evaluate *Mal-ID*.

| Source | Immune state | Size |
| --- | --- | --- |
| Ramesh et al., 2017 (150) | Common variable immunodeficiency | 35 patients, 35 samples<br>916,158 TRB clones |
|  | Healthy, common variable immunodeficiency negative | 22 healthy donors, 22 samples<br>1,229,033 TRB clones |
| Nolan et al., 2020 (79) | Acute Covid-19, sampled between 11 and 21 days after symptom onset with no recorded immunosuppression, cancer, autoimmune disease, or other comorbidities | “NIH/NIAID” cohort: 46 patients, 46 samples<br>2,525,162 TRB clones<br>*Used for training in “held out cohort” experiment* |
|  |  | “ISB” cohort: 9 patients, 9 samples<br>1,235,404 TRB clones<br>*Used for training in “held out cohort” experiment* |
|  |  | “HUniv12Oct” cohort: 33 patients, 33 samples<br>4,403,321 TRB clones<br>*Used for evaluation in “held out cohort” experiment* |
| Towlerton et al., 2022 (151) | HIV, anti-retroviral therapy naive | 30 patients, 60 samples<br>1,222,731 TRB clones |
| Savola et al., 2017 (152) | Sero-positive rheumatoid arthritis, newly diagnosed and treatment naive | 30 patients, 30 samples<br>456,625 TRB clones |
|  | Healthy, rheumatoid arthritis negative | 13 healthy donors, 13 samples<br>246,495 TRB clones |
| Mitchell et al., 2022 (153) | Pediatric type-1 diabetes, new-onset | 143 patients, 143 samples<br>17,114,391 TRB clones |
|  | Healthy, type-1 diabetes negative | 25 healthy donors, 100 samples<br>8,559,072 TRB clones |
| Emerson et al., 2017 (17) | Healthy | Cohort 1: 666 healthy donors, 666 samples<br>90,379,128 TRB clones<br>*Used for training in “held out cohort” experiment* |
|  |  | Cohort 2: 120 healthy donors, 120 samples<br>17,390,850 TRB clones<br>*Used for evaluation in “held out cohort” experiment* |
| immunoSEQ hsTCRB-V4b control data (154) | Healthy | 88 healthy donors, 88 samples<br>8,661,435 TRB clones |

**Table S5:** Model performance for predicting age, sex, and ancestry of healthy individuals with known demographics. The full *Mal-ID* BCR+TCR ensemble architecture was retrained for each task. To cast age as a classification problem, the continuous variable was discretized either into deciles, at a 50-year threshold, or at an 18-year threshold. We report held-out test set performance, averaged over three cross-validation folds, from the metamodel architecture (unregularized logistic regression for the ancestry target, linear support vector machine or ridge logistic regression for the age targets, and random forest for the sex target) with highest AUROC. Abstentions hurt accuracy scores (they count as incorrect predictions) but are not included in the calculation of the probability-based AUROC metric, because no predicted class probabilities are generated for abstained samples. The sex classification is reported for two cross-validation folds, not three as for the other analyses. One cross-validation fold was removed because the CDR3 clustering component (Model 2) abstained on enough of the fold’s validation set that only examples from one class remained for training a metamodel. The sex class imbalance in one fold stems from two design decisions. First, the validation set includes fewer samples than the train or test sets, and it gets even smaller after filtering to healthy donors only for this analysis. Second, we use the same cross-validation splits for all analyses; they were designed to split diseases evenly, not other features like sex.

| Input | Prediction target | Accuracy | AUROC |
| --- | --- | --- | --- |
| BCR+TCR sequence features from 165 healthy samples | Ancestry | 59.4%<br>(9.1% abstentions) | 0.775 |
|  | Age (<20, 20-30, ..., 70-80) | 25.5%<br>(27.9% abstentions) | 0.660 |
|  | Age (under 18, 18 or older) | 54.5%<br>(45.5% abstentions) | 1.0 |
|  | Age (under 50, 50 or older) | 53.9%<br>(19.4% abstentions) | 0.745 |
| BCR+TCR sequence features from 110 healthy samples | Sex | 30.0%<br>(41.8% abstentions) | 0.475<br>(worse than random) |

**Table S6:** Classification results for disease prediction with demographics-aware variants of the *Mal-ID* model. When age is incorporated as a feature, it is treated as a continuous variable. We report held-out test set performance, averaged over three cross-validation folds, from the metamodel architecture with highest AUROC (elastic net logistic regression, random forests, linear support vector machine, lasso logistic regression, elastic net logistic regression, and random forests, respectively). Abstentions hurt accuracy scores (they count as incorrect predictions) but are not included in the calculation of the probability-based AUROC metric, because no predicted class probabilities are generated for abstained samples.

| Input | Prediction target | Accuracy | AUROC |
| --- | --- | --- | --- |
| Age | Disease (358 BCR+TCR samples from individuals with known age, sex, and ancestry) | 28.2% | 0.683 |
| Sex |  | 32.4% | 0.588 |
| Ancestry |  | 65.4% | 0.793 |
| Age, sex, and ancestry |  | 67.9% | 0.851 |
| BCR + TCR sequence features, age, sex, ancestry, and interaction terms between sequence and demographic features |  | 86.3%<br>(1.4% abstentions) | 0.983 |
| BCR + TCR sequences features with age, sex, and ancestry regressed out |  | 81.6%<br>(1.4% abstentions) | 0.963 |

**Table S7:** Validation set performance of different ways to aggregate Model 3 individual sequence predictions into entire repertoire predictions. We report average and standard deviation across three folds for the following strategies: mean, mean trimmed by 5% from the bottom end of the probability distribution, median, and using entropy thresholds to exclude close call sequences (those who have roughly equal predicted probabilities for all classes, i.e. high entropy) from aggregation. An entropy threshold of 1.8 or higher would apply no filtering to six-class predicted probability vectors; we threshold at 80% and 90% of this maximum value to filter out sequences whose predicted class probabilities have high entropy.

| Strategy applied to predicted class probability vectors for all sequences in a sample | BCR AUROC | TCR AUROC |
| --- | --- | --- |
| Mean | **0.881 +/- 0.011** | 0.810 +/- 0.019 |
| Mean, 5% trimmed from bottom | 0.878 +/- 0.013 | 0.809 +/- 0.017 |
| Median | 0.856 +/- 0.014 | 0.785 +/- 0.013 |
| Entropy threshold (90%) | 0.873 +/- 0.016 | 0.896 +/- 0.023 |
| Entropy threshold (80%) | 0.868 +/- 0.008 | **0.912 +/- 0.019** |

**Table S8:** kBET batch effect measurement. Rejection rate of the null hypothesis that the batch distribution in a sequence’s local neighborhood is the same as the global batch distribution, reported as average and standard deviation across 3 folds. Values closer to 0 indicate the null hypothesis is rarely rejected and suggest the batches are well mixed.

| Immune state | BCR | TCR |
| --- | --- | --- |
| Covid-19 | 44.1 +/- 5.7% | 4.8 +/- 0.2% |
| SLE | 18.7 +/- 3.2% | 3.7 +/- 0.3% |
| Healthy/Background | 31.4 +/- 4.5% | 6.6 +/- 0.6% |

